# Orchestration of *Staphylococcus aureus* EV biogenesis by nutrient availability through quorum sensing

**DOI:** 10.64898/2026.04.06.716714

**Authors:** Clariss Limso Yamamoto, Meta Kuehn

## Abstract

The release of extracellular vesicles (EV) is a universally conserved process. Bacterial EVs package diverse cargo, including proteins and nucleic acids, and influence bacterial adaptation and survival as well as host-pathogen interactions. Currently, our understanding of the mechanisms underlying global principles in Gram-positive EV biogenesis and release is limited, partly due to labor-intensive vesicle isolation and assessment methods. Here, we describe a moderately high-throughput approach to analyze the Nebraska Transposon Mutant Library to identify genetic determinants of EV production in *S. aureus*. We show that the *agr* quorum sensing system dictates EV production in response to nutrient availability, likely through communication with the adaptive stress response. This study demonstrates the contribution of nutritional stress to vesiculogenesis and supports a conserved communication strategy that allows metabolic state to influence EV production.

## Introduction

Extracellular vesicles (EV) are membrane-derived proteoliposomal nanoparticles that have been found to be produced in all three domains of life (Deatherage & Cookson, 2012; Gill et al., 2019; Kanehisa et al., 2021). Bacterial EVs are derived from the outermost membrane of the cell and contain a heterogenous mixture of cellular components including lipids, carbohydrates, lipoproteins, soluble and membrane proteins, virulence factors, toxins, nucleic acids, and metabolites. Bacterial EVs have been primarily studied in Gram-negative bacteria, whose EVs are largely derived from the outer membrane, and are aptly named outer membrane vesicles (OMV) (McMillan et al., 2021). OMVs were first observed over 50 years ago through transmission electron microscopy (TEM) studies (Kulp & Kuehn, 2010). Over many years of OMV research, OMVs have been shown to have roles in a variety of bacterial functions, such as horizontal gene transfer, cell-to-cell communication, toxin delivery, biofilm formation and modulation, interactions with host cell, induction of an pro-inflammatory and a protective immune response, among many others (Brown et al., 2015; Ellis & Kuehn, 2010; Kulp & Kuehn, 2010; McMillan et al., 2021).

In comparison, research on EV production by Gram-positive bacteria has lagged, largely due to the decades-long skepticism regarding the ability of EVs to be secreted beyond the thick Gram-positive cell wall (Brown et al., 2015). It was only within the last two decades that EVs from Gram-positive species were isolated and characterized (Lee et al., 2009). Lee and colleagues reported on the proteomic composition of EVs purified from the cultures of Gram-positive bacterium, *Staphylococcus aureus*, confirming the secretion of EVs by Gram-positive bacteria. Subsequent studies have now identified EV production to occur in various Gram-positive bacteria (Briaud & Carroll, 2020; Brown et al., 2015; Liu et al., 2018).

Analyses of the contents of Gram-positive derived EVs have shown, similar to OMVs, a heterogeneous mixture of cellular components originating from its parent cell (Bose et al., 2020; Briaud & Carroll, 2020; Brown et al., 2015; McMillan et al., 2021). Proteomic analyses of EVs from various Gram-positive bacteria revealed that cytoplasmic proteins, virulence factors, and toxins, comprise a large portion of the Gram-positive EV protein cargo. Toxins have been identified in the EVs of various Gram-positive bacteria including *S. aureus*, *Streptococcus pneumoniae*, *Streptococcus pyogenes*, *Streptococcus agalactiae*, and *Bacillus anthracis*, to name a few. Nutrient-scavenging components have also been identified in Gram-positive EVs. For example, iron-binding proteins that aid in iron storage and survival in iron-depleted conditions have been identified in *S. aureus, Dietzia* sp. DQ12-45-1b, as well as *Streptomyces coelicolor* EVs (Askarian et al., 2018; Gurung et al., 2011; Lee et al., 2009; Schrempf et al., 2011; M. Wang et al., 2021; Wang et al., 2018). It is still unknown to what degree protein cargo are sorted and loaded into EVs, however enrichment and depletion of specific proteins in EVs compared to the parent cell have been observed, suggesting the existence of sorting mechanisms that package EV cargo (Briaud & Carroll, 2020; Jeon et al., 2017; Resch et al., 2016; Tartaglia et al., 2020).

For example, in *S. aureus*, a study comparing EVs from *S. aureus* strains originating from diverse hosts identified a highly conserved core *S. aureus* EV proteome (Tartaglia et al., 2020). In addition to protein cargo, selective sorting of lipids and an nucleic acids into Gram-positive EVs have also been observed, strongly suggesting a conserved EV cargo sorting mechanism (da Luz et al., 2021; Jeon et al., 2018; Joshi et al., 2021; Resch et al., 2016).

The functions of EVs are largely dependent on their cargo, and the presence of toxins, siderophores, immune evasion proteins, adhesins, and antibiotic resistance proteins clearly indicate a role for EVs in virulence. In *S. aureus*, several reports have demonstrated the immunostimulatory properties of EVs on mammalian host cells (Hong et al., 2014; Rodriguez & Kuehn, 2020; Tartaglia et al., 2018; Wang et al., 2020). It has also been suggested that EV-associated toxins are more potent than the soluble secreted form of the same toxin. In *S. aureus*, EV associated α-hemolysin, but not soluble α-hemolysin, triggered keratinocyte necrosis in keratinocytes (Hong et al., 2014).

Although some aspects of Gram-positive EVs have been explored and uncovered, major outstanding questions in the field concern EV biogenesis and regulation of their production. A previous study described a biogenesis and release pathway for EVs in *S. aureus* dependent on the Staphylococcal alpha-type-phenol-soluble modulins 1-4 (α-PSM1, α-PSM2, α-PSM3, and α-PSM4), which we will refer to here as α-PSMs collectively (Wang et al., 2018). α-PSMs are small, amphipathic, α-helical peptides that act as virulence determinants and contribute to *S. aureus* pathogenesis (Cheung et al., 2014; Peschel & Otto, 2013; Wang et al., 2007). Wang and colleagues showed that α-PSMs can promote vesicle formation presumably by disruption of the cytoplasmic membrane due to their surfactant-like activity. Schlatterer et al. later showed that α-PSMs promote vesicle formation by increasing membrane fluidity (Schlatterer et al., 2018).

Wang et al. also showed that the degree of peptidoglycan crosslinking is inversely proportional to EV yield (Wang et al., 2018). EV production and EV size increased for 1) *S. aureus* treated with Penicillin G, which decreases peptidoglycan crosslinking; 2) the *S. aureus* mutant lacking penicillin-binding protein 4 (PBP4), a carboxypeptidase that is essential for secondary crosslinking of the peptidoglycan; and 3) the *S. aureus* mutant lacking the gene *tagO*, encoding the enzyme catalyzing the first step of wall teichoic acid biosynthesis. Conversely, a decrease in EV yield was found for a mutant lacking the gene encoding Sle1, an autolysin that belongs to a family of peptidoglycan hydrolases that play a role in the separation of daughter cells. A separate study has also demonstrated the effect of peptidoglycan crosslinking on EV formation: Andreoni and colleagues found that β-lactam antibiotics, which increase the permeability of peptidoglycan, can induce the formation of EVs (Andreoni et al., 2019).

In addition to the prophage-independent mechanisms of EV biogenesis described above, a relationship between EV production and phage induction has also been reported. Andreoni and colleagues demonstrated that oxidative stress-inducing antibiotics can promote *S. aureus* EV production through induction of phage (Andreoni et al., 2019). Induction of the SOS-response, which results in phage induction in lysogenic *S. aureus* strains, resulted in an increase in EV formation only in lysogenic *S. aureus* strains but not in non-lysogenic strains. Along with the induction of EV formation in the lysogenic strains, they observed the presence of ghost cells, suggesting that the death of the mother cell is linked to phage-dependent EV formation. Wang et al. also investigated the role of phages in EV formation (Wang et al., 2018). The *S. aureus* strain NCTC 8325 carrying ϕ11, ϕ12, and ϕ13 was demonstrated to induce phages in the conditions they used, while the strain 8325-4, which is cured of all three prophages, did not. They further showed that the lysogenic and non-lysogenic strains produced comparable EV amounts. Thus, it remains unclear whether the phage-dependent EV formation mechanism described by Andreoni and colleagues is a strain-specific phenomenon, or is only activated in certain conditions, such as the activation of the SOS-response pathway.

A virulence-associated genetic pathway has also been implicated in *S. aureus* EV production. *S. aureus* mutants lacking AgrA were reported to yield undetectable levels of EVs (Im et al., 2017). AgrA is a component of the quorum-sensing accessory gene regulator (*agr*) system, which controls a large arsenal of virulence factors including α-PSMs. AgrA, specifically upregulates *α-psm* transcription by binding directly to its promoter domain (Thoendel et al., 2011). Recently, delta-hemolysin (*hld*), whose transcript is embedded within the *agr*-regulated small RNA, RNAIII, was demonstrated to modulate vesiculogenesis and influence the properties of *S. aureus* EVs (Chen et al., 2023; Wang et al., 2023). Thus, it is likely that EV formation in *S. aureus* is, at least in part, regulated by the *agr* system.

The general stress response has also been linked to EV production. A point mutation in the alternative sigma factor B (σ^B^), which modulates stress responses in several Gram-positive bacteria, resulted in an increase in vesicle production in *S. aureus* (Qiao et al., 2022; van Schaik & Abee, 2005). β-galactosidase reporter and gel-shift assays demonstrated that the σ^B^ point mutation weakened the ability of σ^B^ to bind the *nuc* gene promoter, resulting in significant reduction in the expression of the secreted nuclease (Nuc) (Qiao et al., 2022). The *nuc* deletion mutant showed an increase in vesicle production, and functional complementation restored vesicle production to wild-type levels. A follow up study found that the σ^B^ point mutation also hindered the ability of σ^B^ to bind to the *asp23* gene promoter, resulting in the decreased expression of the alkaline shock protein 23 (Asp23), and consequently increased vesicle production (Li et al., 2024). Similarly, another group found that addition of the compound, rhodomyrtone, reduced σ^B^ activity in *S. aureus* during exponential growth and resulted in significantly decreased vesicle production (Mitsuwan et al., 2019).Accordingly, it seems that the general stress response and the stress response transcription regulator, σ^B^, have regulatory roles in EV biogenesis.

Both σ^B^ and AgrA are global regulators that control the expression of numerous genes (Thoendel et al., 2011). Consequently, their involvement in the production of EVs suggests that vesiculogenesis is not dependent on a small number of genes - Instead, a complex global gene network may be involved.

In this study, we aimed to broadly identify genetic determinants of EV production in Gram-positive bacteria using an unbiased phenotypic screening approach. Traditional methods of vesicle isolation to assess bacterial EV production are laborious and time consuming, hindering high-throughput global approaches, such as the analysis of transposon mutant libraries (Nasukawa et al., 2021; Prados-Rosales et al., 2014). Previously, a moderately high-throughput method to assess OMV production by Gram-negative bacteria was successfully conducted to screen *E. col* transposon mutants and the non-essential Keio mutant strain library, yielding novel insights into OMV biogenesis and regulation (Kulp et al., 2015; McBroom et al., 2006). Here, we describe the development of a method to assess vesicle production in all non-essential mutant *S. aureus* strains in the Nebraska Transposon Mutant Library (NTML) (Fey et al., 2013). We demonstrate that this moderately high-throughput screen can successfully identify mutants with vesiculation phenotypes by verification using the traditional ultracentrifugation method of EV isolation. We also show that EVs can aid in adaptation and survival during stress. Additionally, our results demonstrate that *S. aureus* EV biogenesis is largely orchestrated though the *agr* quorum sensing system, and may be, in part, upregulated in response to nutrient limitation. Lastly, our results suggest that EV biogenesis may be governed by crosstalk between several cellular processes including the quorum sensing system and the stringent response, suggesting a possible universally conserved communication strategy that allows metabolic state to influence vesiculogenesis.

## Results

### Vesiculation Screen Development

With this study, we sought to gain global, unbiased insight into the genetic basis for the mechanism and regulation of EV production in *S. aureus*. The existing methods to measure Gram-positive bacterial vesiculation levels are based on large-scale, and time-consuming assays that are not suitable for genome-wide studies of vesiculation phenotypes (Nasukawa et al., 2021; Prados-Rosales et al., 2014). However previous success based on a 96-well assay to detect vesiculation phenotypes of *E. coli* mutant libraries revealed genes critical for OMV biogenesis and regulation for Gram-negative bacteria (Kulp et al., 2015; McBroom et al., 2006). Therefore, we designed experimental and analytical methods to assess vesicle production in small volume *S. aureus* cultures in a moderately high-throughput, cost-effective, and reproducible fashion.

Key parameters needed to be established in the initial design of the screen: the timing of phenotype assessment during culture growth, and the identification of mutants that have bacterial integrity defects in the culture conditions. To decide on a timepoint for the screen we relied on two considerations: first, late exponential phase cultures exhibit high viability and have had time to produce a quantifiable amount of EVs; and second, prior studies have shown that EV production in *S. aureus* is partly regulated by the *agr* quorum sensing system (Im et al., 2017). Because *agr* is a quorum-sensing regulon, maximal *agr* activity occurs during exponential growth, and is followed by a sharp drop during stationary phase (Geisinger et al., 2012). Therefore, we chose late exponential/early stationary phase as a timepoint and needed to determine when cultures grown in the screening conditions would attain this phase. A 1:40 dilution of *S. aureus* JE2 overnight culture into fresh tryptic soy broth (TSB) in 96-well microtiter plates result in very reproducible growth, with the late exponential/early stationary phase consistently occurring at around five hours of growth (Figure 1A). Thus, we chose to assess vesicle production of *S. aureus* cultures after five hours of growth. In order to rule out apparent high vesiculation values due to cell lysis, a separate assay to determine the membrane integrity of cells was necessary. Previously, a rapid 96-well assay, more sensitive than the commercially available LIVE/DEAD BacLight assay, was developed for assessing viability of *Borrelia burgdorferi* in a high throughput format (Feng et al., 2014). In this assay, the ratio of viable and compromised cells is determined using membrane permeant (SYBR Green) and non-permeant (propidium iodide, PI) nucleic acid dyes. We compared the conventional LIVE/DEAD BacLight assay to a modified SYBR Green / PI viability assay, in which SYBR Green was substituted with a similar nucleic acid dye, SYBR Gold. *S. aureus* JE2 was grown to early stationary phase, and the culture was assigned to represent a 100% viable culture. An aliquot of the 100% viable culture was heat killed by boiling for 20 minutes and assigned to represent a 0% viable culture. The 100% and 0% viable cultures were then mixed in various ratios, diluted by 2 and 10-fold before being assessed for viability using either the LIVE/DEAD BacLight assay or the modified SYBR Gold/PI assay. In the samples diluted 2-fold, both viability assays showed a linear relationship between the signal and ratio of live to dead cells, with the R^2^ values of the lines of best fit being 0.99 for the LIVE/DEAD BacLight assay and 0.94 for the SYBR Gold/PI assay (Figure 1B). However, the LIVE/DEAD BacLight assay lost sensitivity when the samples were diluted 10-fold (R^2^=0.86, Figure 1B). On the other hand, the SYBR Gold/PI viability assay retained its sensitivity even in the 10-fold diluted samples (R^2^=0.97). We therefore chose the SYBR Gold/PI viability assay to identify mutants with compromised membrane integrity and exclude them from the screen results.

**Figure 1:**
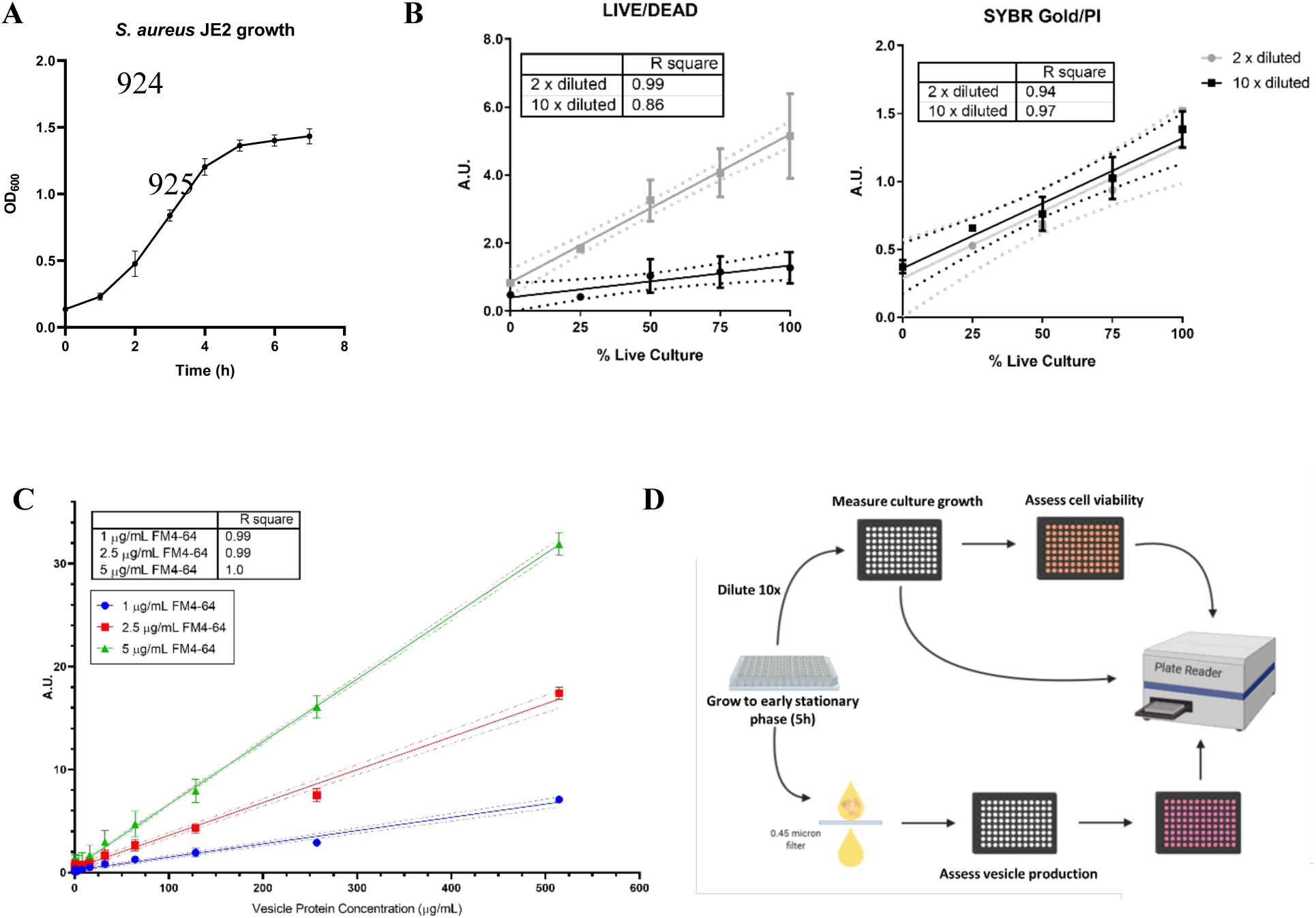
Development of the moderately high-throughput vesiculation screen. **A.)** *S. aureus* JE2 growth curve in 96-well microtiter plates graphed as averages from three independent experiments expressed as mean ± standard deviation **B.)** Comparison of LIVE/DEAD assay with SYBR Gold/PI viability assay in 2-fold or 10-fold diluted mixture of cultures with varying ratios of live and dead cell cultures. Averages from three independent experiments are presented as mean of LIVE/DEAD ratios or SYBR Gold/PI ratios ± standard deviation. Fluorescence was monitored by excitation at 488nm and taking emission values at 500nm for SYTO9 (LIVE) and 635nm for propidium iodide (PI; DEAD) and 535nm for SYBR Gold. Least squares regression analysis was performed and dashed lines represent the 95% confidence bands for the best fit line. R^2^ values were determined as a measure goodness of fit. **C.)** Determination of FM4-64 sensitivity to detect EVs. Purified *S. aureus* JE2 EVs were diluted with TSB to varying concentrations as determined by protein content. 1μg/ml, 2.5μg/ml, and 5μg/ml concentrations of FM4-64 were used to detect lipid content of EVs. FM4-64 fluorescence was measured by excitation at 506 nm and reading emission values at 750nm. Averages from three independent experiments are presented as mean ± standard deviation. Least squares regression analysis was performed and dashed lines represent the 95% confidence bands for the best fit line. R^2^ values were determined as a measure goodness of fit. **D.)** Graphic summary of developed screen. Overnight NTML mutants were diluted 1:40 and grown for 5 hours to early stationary phase. Cell culture growth was measured by OD_600_, cell viability by SYBR Gold/PI viability assay, and vesiculation by FM4-64.

In order to assess EV production, the culture medium containing secreted extracellular materials, including EVs, were sterile-filtered through 96-well, 0.45μm, PVDF, filter plates. The lipophilic dye, FM4-64, was then used to quantify EVs in the filtered medium. FM4-64 exhibits low fluorescence in aqueous environments but fluoresces intensely in a lipophilic environment and is regularly used to stain EVs for quantification of lipid content. 1, 2.5, and 5 µg/ml concentrations of FM4-64 were tested on a range (0.25 µg/ml to 514.5 µg/ml by protein content) of dilutions of purified vesicles from *S. aureus* (Figure 1C). The FM4-64 values observed in the filtered spent media of the wild-type JE2 strain typically fall between 3 to 8 a.u., thus both 2.5 and 5 µg/ml FM4-64 can be used to effectively quantify EV production (Supplemental Figure 1). For a more cost-effective screen design, 2.5 µg/ml FM4-64 was selected for the screen. To adjust vesiculation values to account for any differences in growth, the FM4-64 signal was normalized to the final optical density (OD_600_) of the culture.

Figure 1D summarizes the overall process of the screen developed. Overnight cultures of mutant strains were diluted 1:40 (5 µl culture into 200µl TSB) with pre-warmed TSB and grown at 37°C with shaking (250 rpm) for 5 hours to late exponential phase in 96-well microtiter plates placed within a humidity chamber to prevent evaporation. A culture aliquot (20 µl) was taken and diluted 10-fold to measure OD_600_ and cell viability by the SYBR Gold / PI viability assay. The remaining culture (160 µl) was filtered through 0.45μm PVDF filter, and 2.5 µg/ml FM4-64 was used to quantify EV production.

Mutants were considered to have significant hypo- or hyper-vesiculation phenotypes if: 1) their cell viability value (SYBR Gold/PI) was more than 2-mean average deviations (MAD) below the mean viability value of the strains within the plate, and 2) their vesiculation value (FM4-64) normalized to their growth (OD_600_) was outside 2-MADs from the mean normalized vesiculation value (FM4-64/OD_600_) of the strains within the plate.

### Pathway analysis of screen results

A total of 173 mutants, 100 hypervesiculating and 73 hypovesiculating, met the defined criteria for having vesiculation phenotypes. We note that 80 mutants were excluded from EV phenotype evaluations due to low viability (Tables 1 and 2, Supplementary Table 1).

**Table 1:**
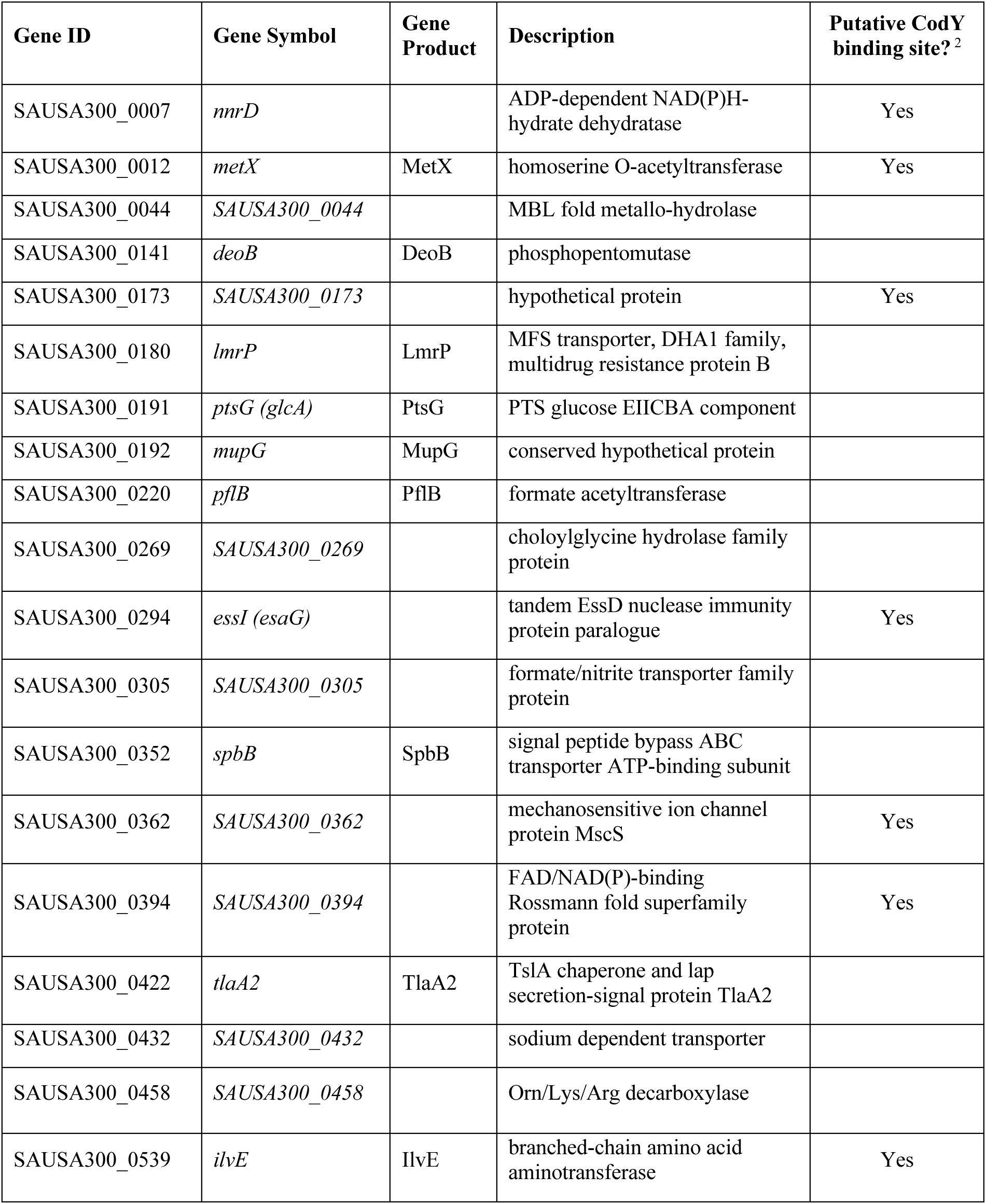
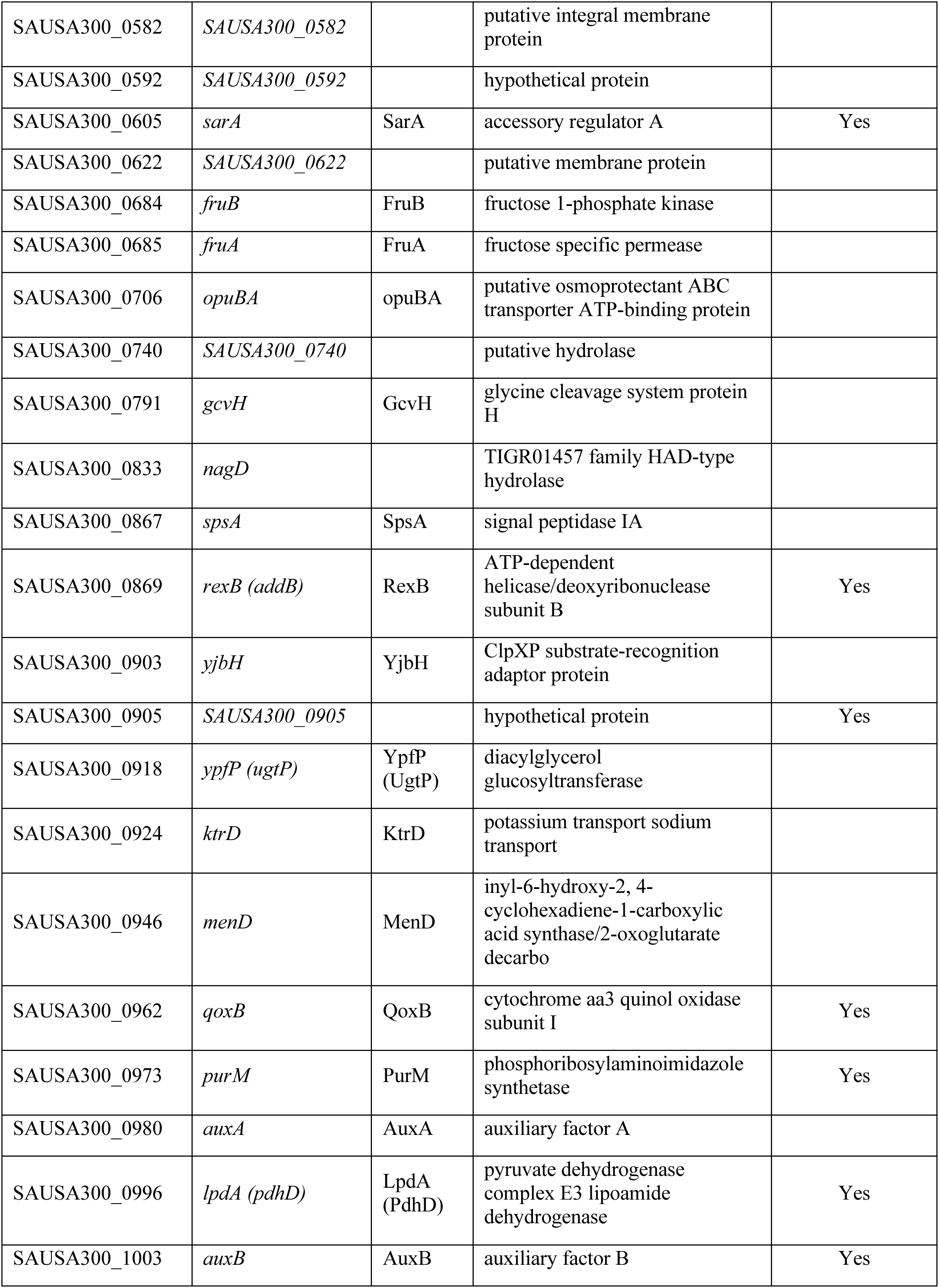
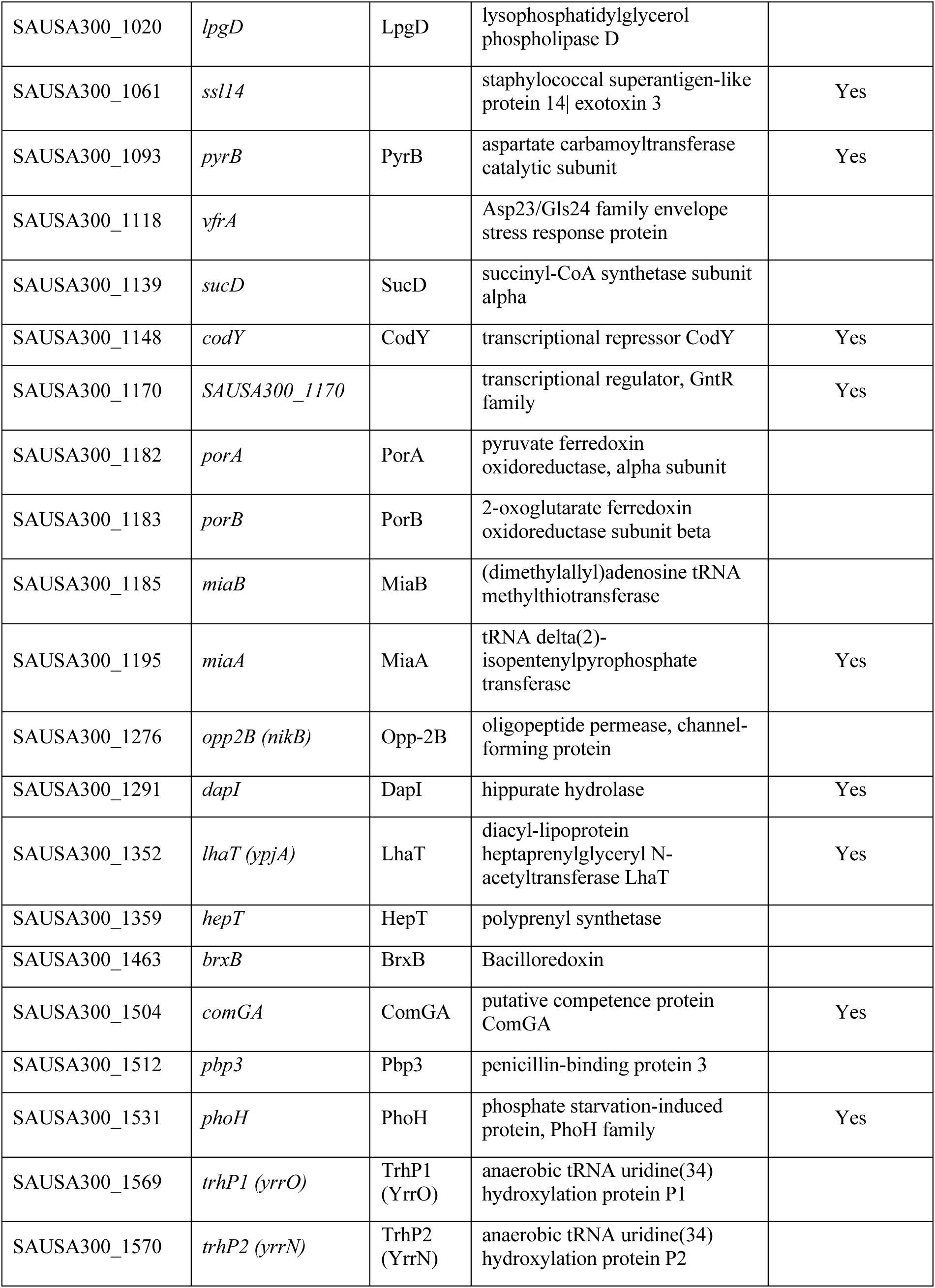
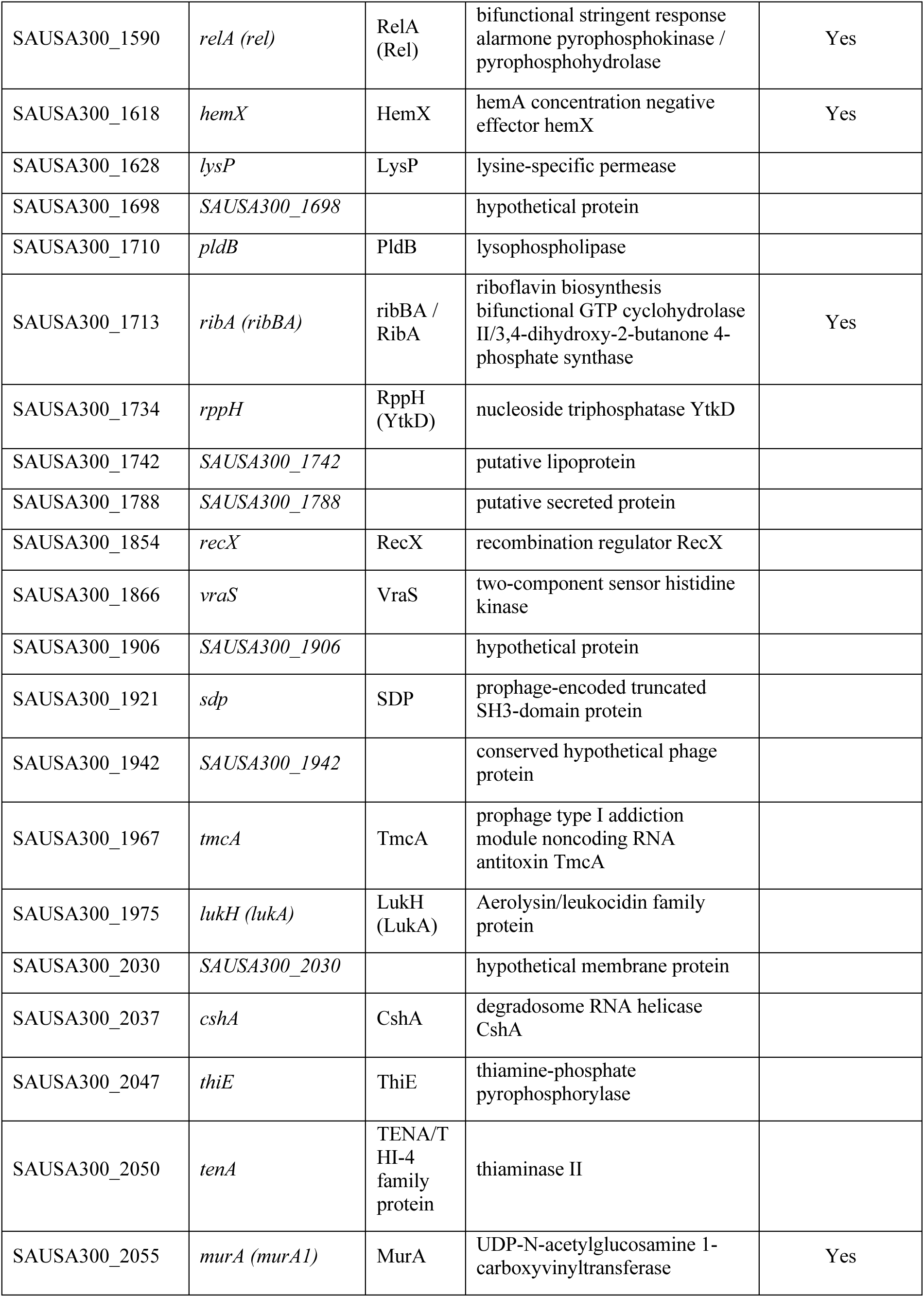
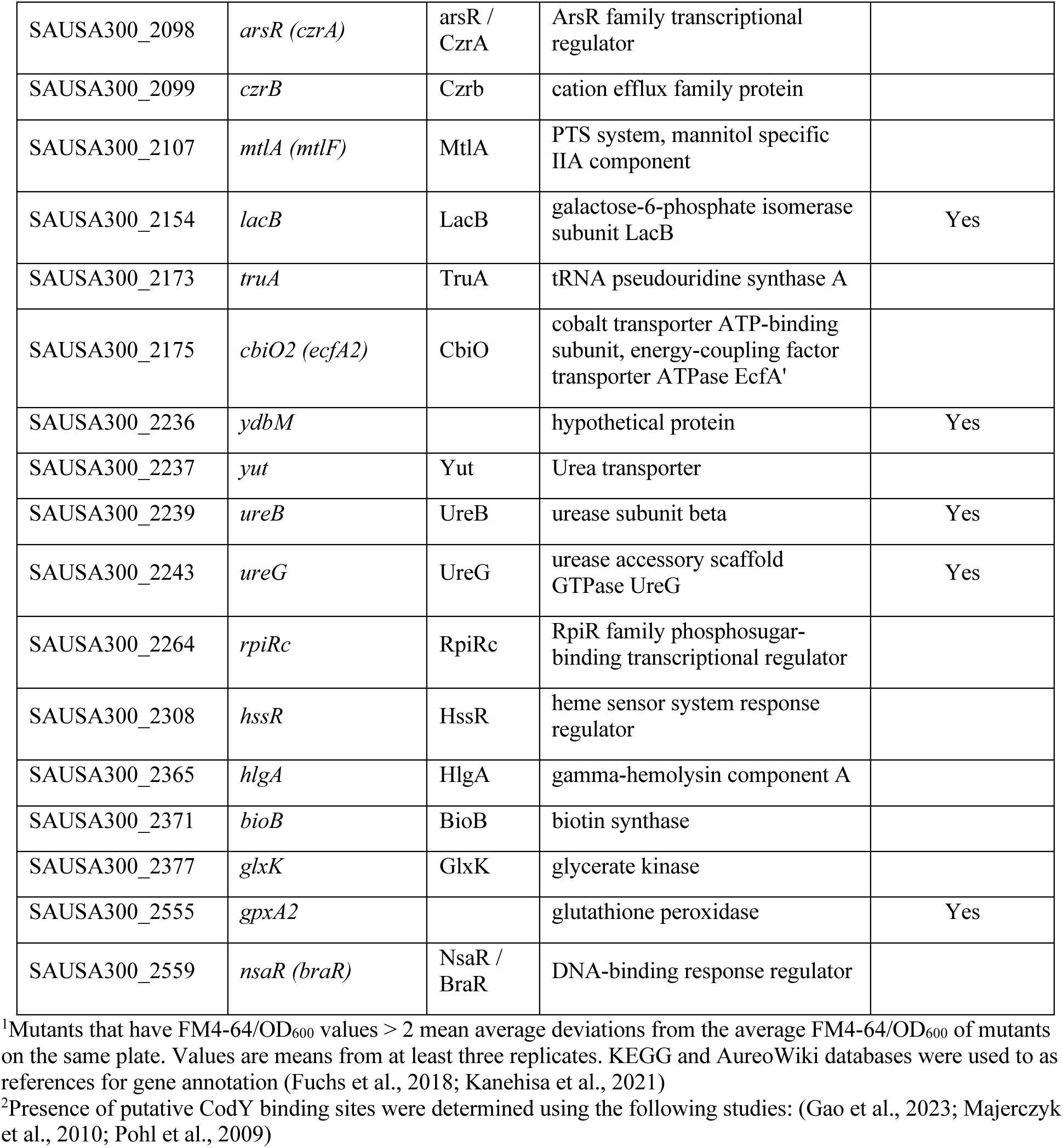
Hypervesiculating *S. aureus* mutants identified in the screen^1^.

**Table 2:**
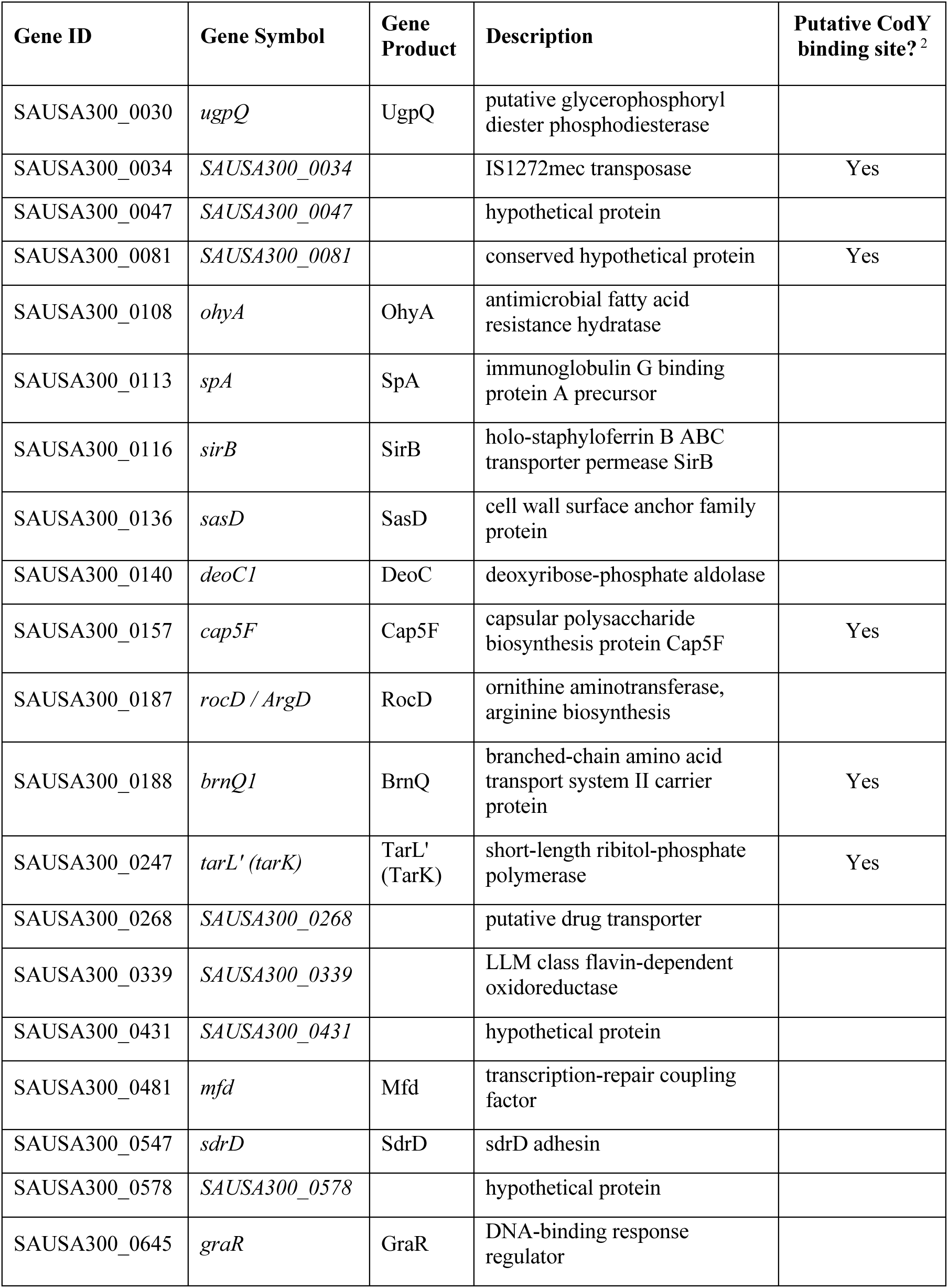
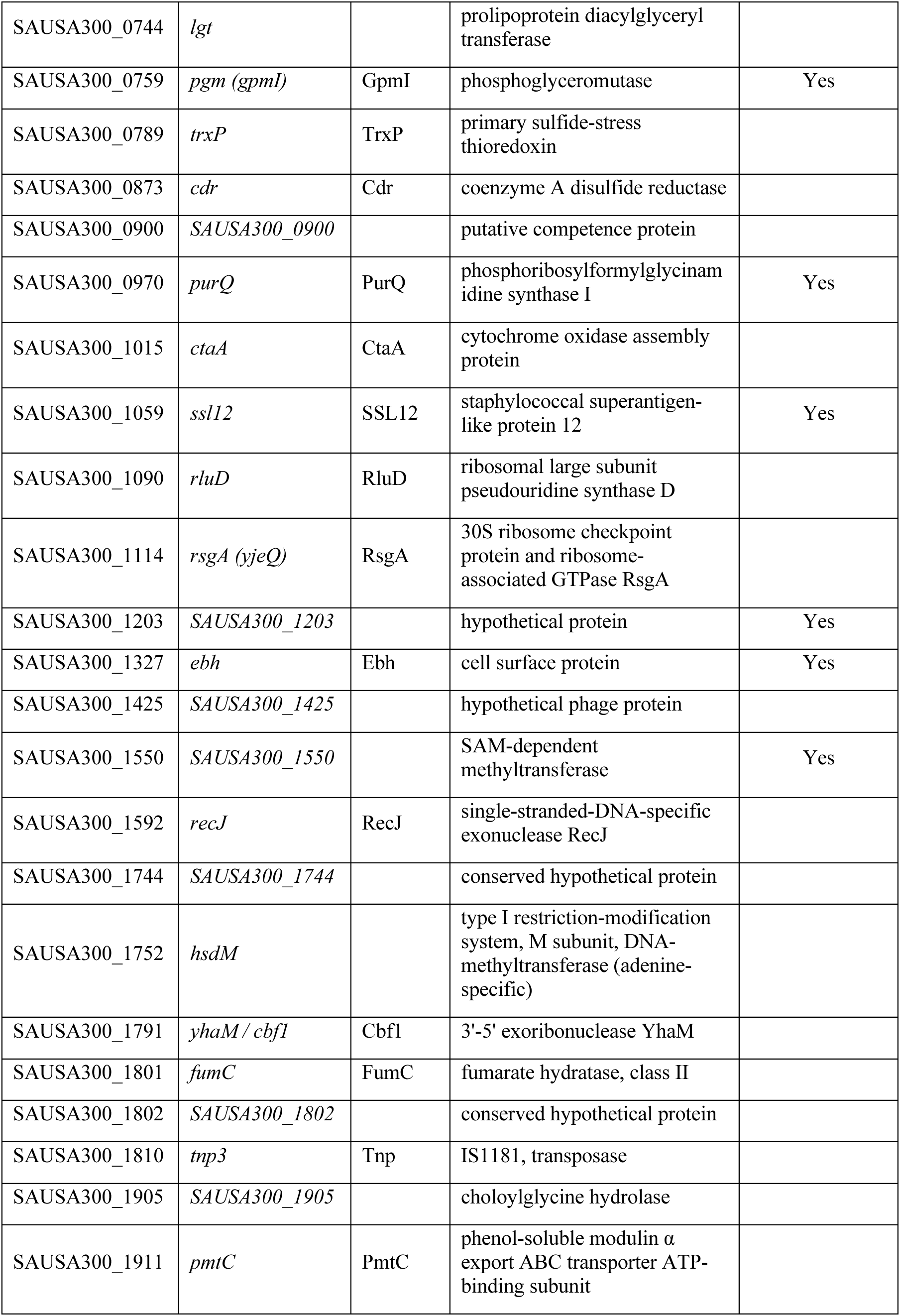
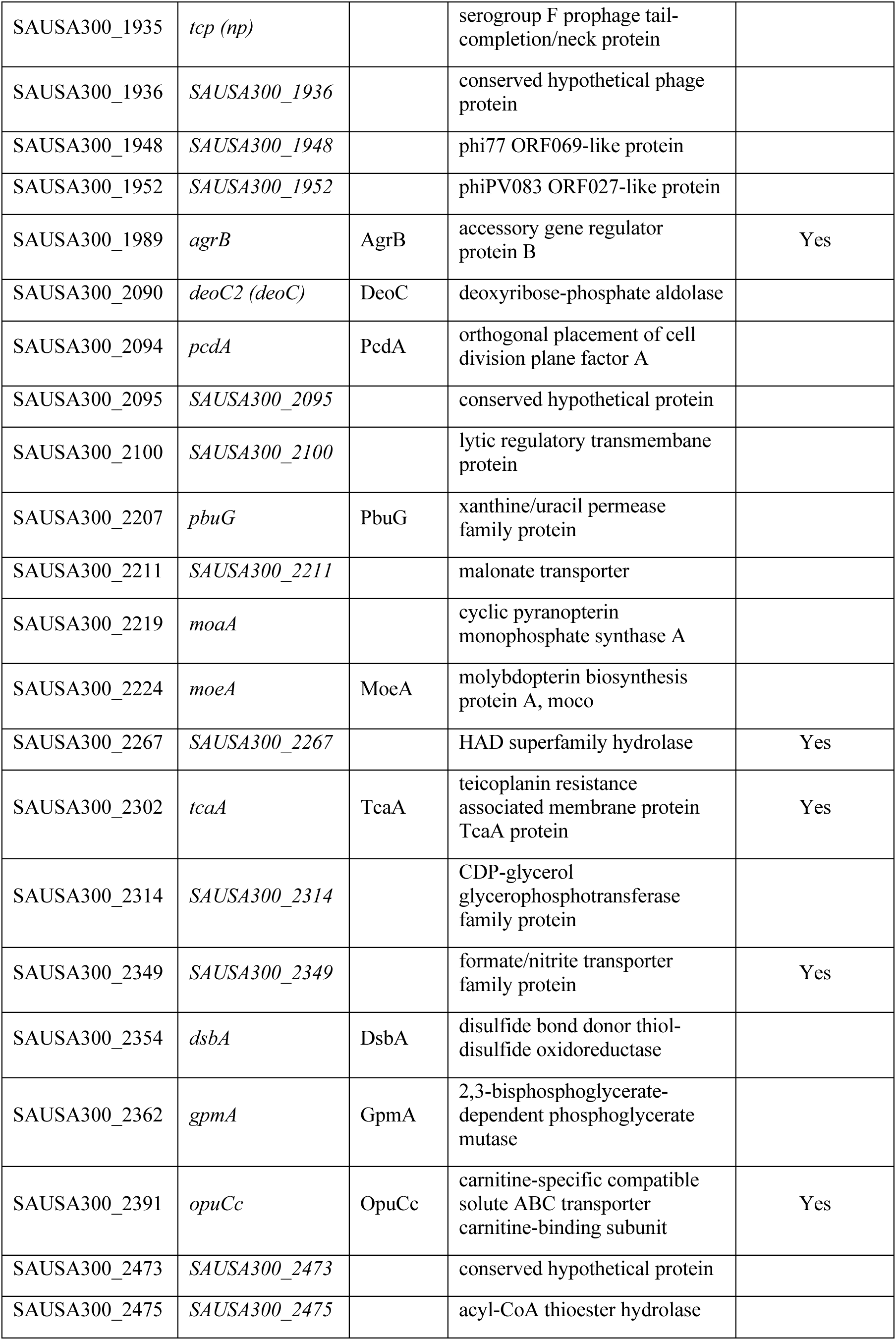
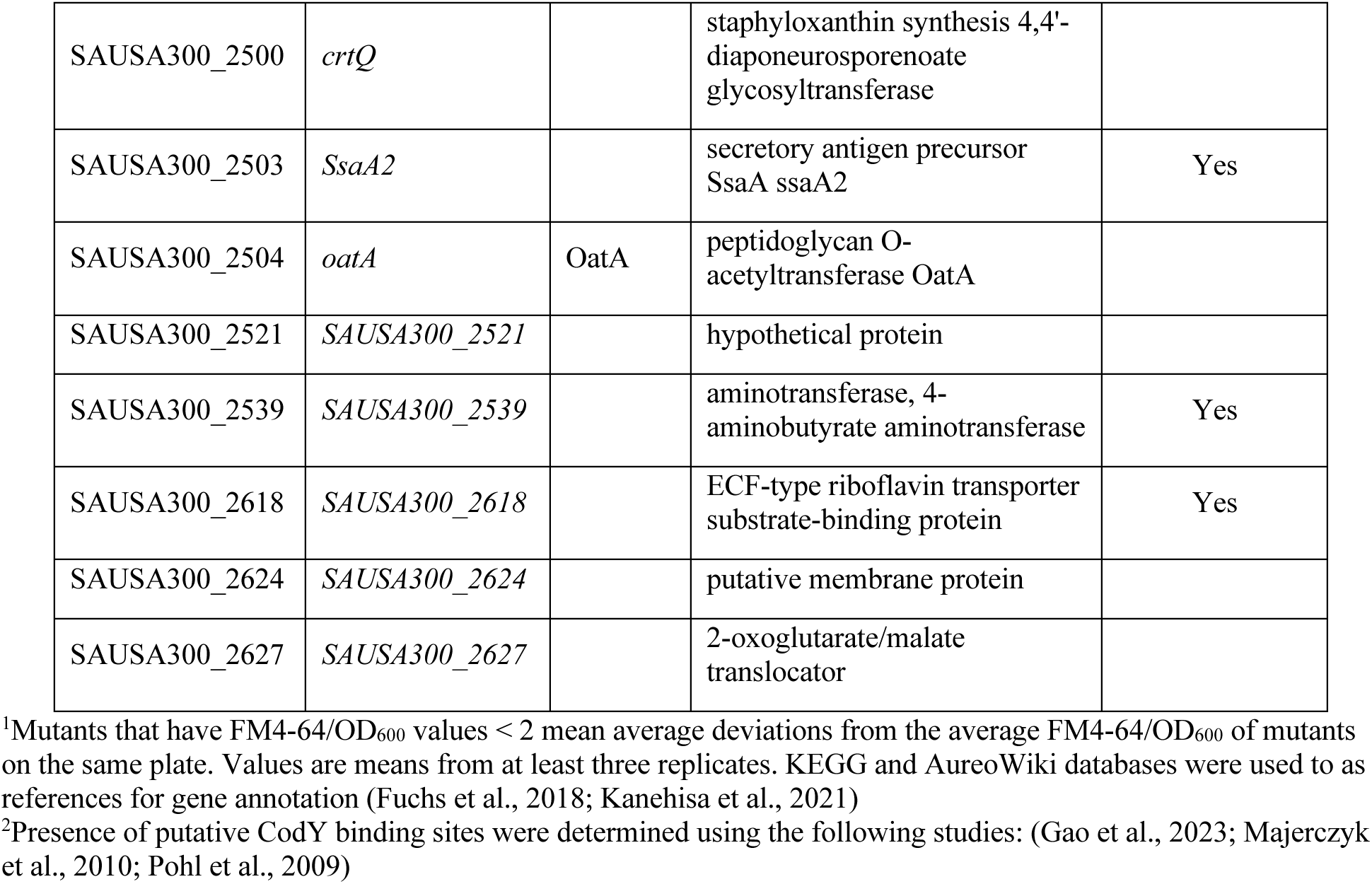
Hypovesiculating *S. aureus* mutants identified in the screen^1^.

We next performed a Gene Ontology (GO) analysis to determine if specific cellular processes or pathways were enriched in our dataset. Because the GO database is not annotated for *S. aureus* JE2 or any other USA300 strain, the strain NCTC8325 was used as the reference strain. Of the 173 gene mutations identified to cause vesiculation phenotypes in the JE2 strain, only 161 are present in the strain NCTC8325. Therefore, the 13 missing genes were excluded from the GO analysis. Many of the identified mutants in the screen were disrupted in genes that code for hypothetical proteins or proteins of unknown functions. Of the genes that had assigned functions, the analysis identified the enrichment of genes involved in cell wall biogenesis, as expected based on prior studies, in the set of identified mutants with vesiculation phenotypes (Table 3, Supplementary Table 2 for full table). In addition to the previously identified pathways and processes that affect *S. aureus* vesiculogenesis, the GO analysis also showed the vesiculation mutant dataset to have an enrichment of genes involved in stress responses, including transcription factors and two-component systems that control the expression of numerous genes in response to stimuli; as well as genes involved in altering membrane fluidity, including branched chain amino acid (BCAA; isoleucine, leucine, and valine) biosynthesis and lipid biogenesis genes, among others.

**Table 3:** GO Analysis^1^.

| **Pathway<sup>1</sup>** | **nGenes** | **Pathway Genes** | **Fold Enrichment** |
| --- | --- | --- | --- |
| Hexose catabolic process | 2 | 5 | 20.20 |
| Transcriptional regulatory protein, C terminal | 3 | 8 | 11.65 |
| Nucleotide catabolic process | 4 | 12 | 10.35 |
| Hexose metabolic process | 4 | 20 | 10.10 |
| Glucose metabolic process | 3 | 18 | 8.42 |
| Teichoic acid biosynthetic process | 3 | 19 | 7.97 |
| Aerobic respiration | 3 | 22 | 6.89 |
| Vitamin biosynthetic process | 4 | 32 | 6.31 |
| GTP-binding | 5 | 25 | 6.21 |
| Signaling | 5 | 41 | 6.16 |
| Guanyl nucleotide binding | 6 | 31 | 6.01 |
| Mostly uncharacterized, incl. Biosynthesis of various secondary metabolites - part 3, and DNA binding HTH domain, AraC-type | 6 | 34 | 5.48 |
| Carbohydrate derivative catabolic process | 5 | 29 | 5.35 |
| Citrate cycle (TCA cycle), and Branched-chain amino acid metabolic process | 6 | 37 | 5.04 |
| Mixed, incl. Biosynthesis of various secondary metabolites - part 3, and Two-component system | 7 | 45 | 4.83 |
| Peptidoglycan-based cell wall biogenesis | 4 | 46 | 4.39 |
| Ion transmembrane transport | 8 | 94 | 4.30 |
| RNA modification | 4 | 47 | 4.30 |
| Cell wall biogenesis | 4 | 47 | 4.30 |
| Cell wall organization or biogenesis | 5 | 61 | 4.14 |
| Lipid metabolic process | 6 | 75 | 4.04 |
| RNA processing | 5 | 65 | 3.89 |
| Cellular lipid metabolic process | 5 | 66 | 3.83 |
| Nucleoside monophosphate biosynthetic process, and Nucleobase-containing compound catabolic process | 7 | 59 | 3.68 |
| Response to stimulus | 13 | 183 | 3.59 |
| Regulation of transcription, DNA-templated | 9 | 129 | 3.52 |
| Cellular response to stimulus | 9 | 132 | 3.44 |
| Response to stress | 7 | 103 | 3.43 |
| Regulation of cellular metabolic process | 10 | 157 | 3.22 |
| Winged helix-like DNA-binding domain superfamily | 9 | 91 | 3.07 |
| Nucleoside phosphate metabolic process | 11 | 112 | 3.05 |
| Transmembrane transport | 22 | 291 | 2.35 |
| Purine ribonucleoside triphosphate binding | 20 | 311 | 2.00 |

<sup>1</sup>GO Analysis was performed using the ShinyGO version 0.85 database (Ge et al., 2020). KEGG and AureoWiki databases were used to as references for gene annotation (Fuchs et al., 2018; Kanehisa et al., 2021). The strain NCTC8325 was used as the reference strain. (Supplementary Table 3 for full table)

### Regulation of EV production by CodY

One of the hypervesiculating mutants we identified is the *codY* mutant. CodY is a highly conserved global transcriptional repressor for genes involved in the adaptive response to BCAA starvation (Geiger & Wolz, 2014; Kaiser & Heinrichs, 2018; Pohl et al., 2009). CodY senses nutrient availability by direct interaction with metabolite effectors, GTP and BCAAs. The CodY regulon includes the genes that code for the enzymes involved in the biosynthesis and metabolism of BCAAs, as well as other amino acids namely, cysteine, glutamate, histidine, serine, tryptophan, tyrosine, aspartate, lysine, threonine, and methionine. CodY also regulates the expression of genes involved in the *agr* quorum sensing system, central metabolism and carbon flow, nitrogen assimilation, various transport systems, as well as virulence genes. Interestingly, we observed a large number of CodY targets in our vesiculation screen dataset (Tables 1 and 2). Of the 173 genes identified to alter vesiculation, 51 (29.5%) also harbor the CodY consensus binding motif within their intergenic and/or coding regions (Gao et al., 2023; Majerczyk et al., 2010; Pohl et al., 2009).

In order to explore the possible role of CodY in EV biogenesis, as well as to validate our moderately high throughput screen data, a larger-volume, quantitative assay using the conventional ultracentrifugation-based vesicle isolation protocol was performed on several mutants including the *codY* mutant, as well as mutants of both CodY-dependent and - independent genes. Vesicle production normalized by OD_600_, was assessed by lipid and protein measurements by FM4-64 and Bradford assays respectively, and compared to that of the wild type parent strain, JE2. Figure 2A demonstrates that the moderately high-throughput vesiculation screen developed in this study successfully identified the vesiculation phenotypes for all the selected mutants we tested. Intriguingly, we observed that the changes in the lipid content are not always proportional to the changes in the protein content of the vesicles produced by the mutants, suggesting that perhaps cargo loading and vesicle budding are regulated independently of each other.

**Figure 2:**
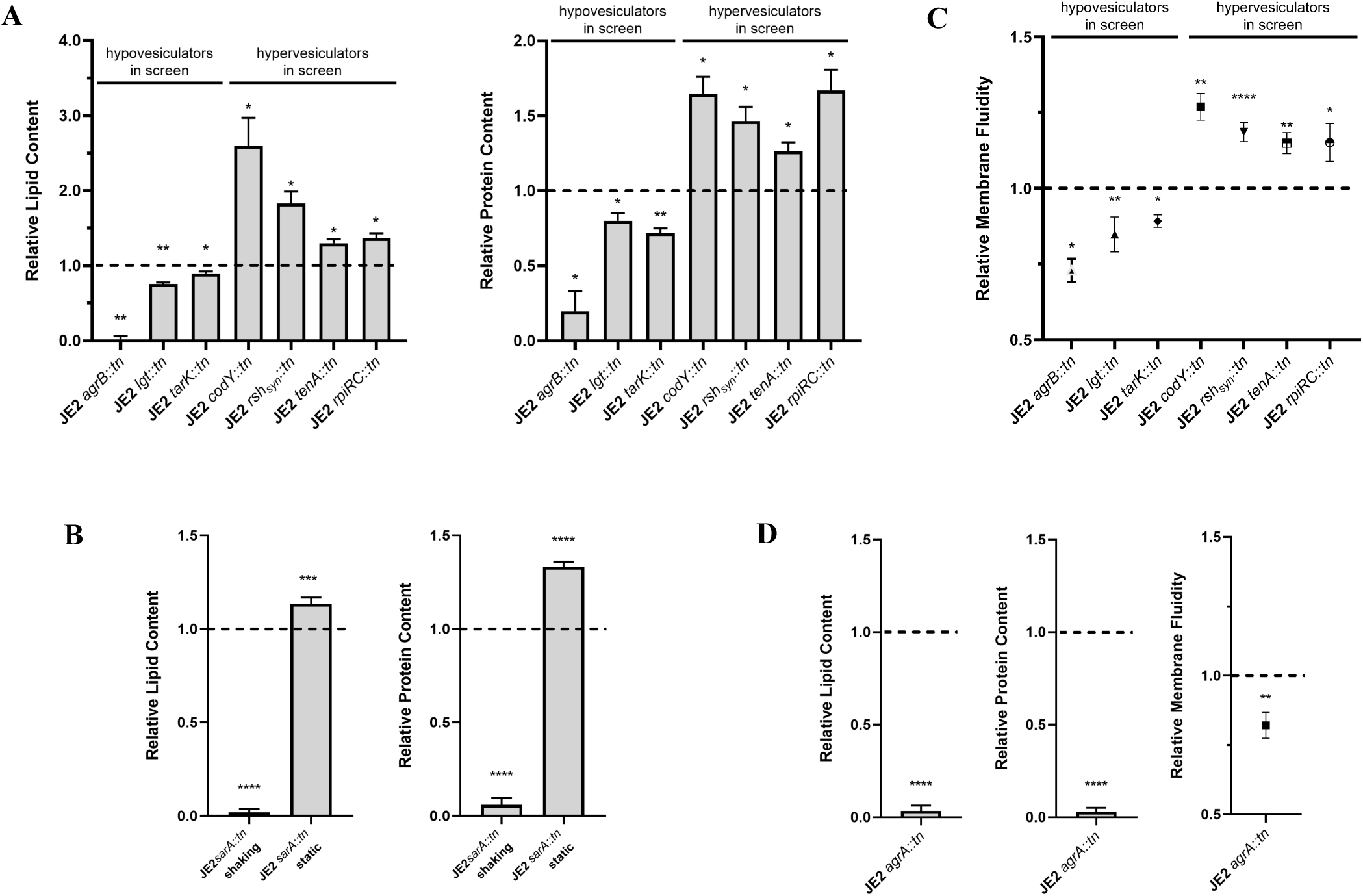
Assessing vesiculation phenotype and membrane fluidity of mutants. **A.)** Lipid and protein contents of EVs purified from vesiculation mutants were assessed by FM4-64 and Bradford Assay respectively and compared to that of the wild type JE2 parent strain represented by the dashed lines. **B.)** The *sarA::tn* mutant’s vesiculation phenotype was inconsistent with the screen result when cells were grown in a flask with shaking. Static growth in a beaker phenocopy the screen result. Lipid and protein contents of purified EVs from were assessed by FM4-64 and Bradford Assay respectively and compared to that of the wild type JE2 parent strain grown in a flask with shaking, represented by the dashed lines. **C.)** Relative membrane fluidity of *S. aureus* cells were measured using pyrenedecanoic acid (PDA). PDA fluorescence was monitored by excitation at 360nm and taking emission values at 400nm (monomer) and 470nm (excimer). Relative membrane fluidity was calculated as a ratio of excimer to monomer fluorescence and normalized to that of the wild type JE2 parent strain represented by the dashed line. **D.)** Vesiculation and relative membrane fluidity of JE2 *agrA::tn* mutant. For all Figures, values are presented as means from three independent experiments ± SD normalized to the control. Samples were compared using mixed effects analysis with Dunnett’s multiple comparison test (**A** and **C**), one-way ANOVA with Dunnett’s multiple comparison test (**B**), or an unpaired t-test (**D**) *p< 0.05, **p< 0.01, ***p< 0.001, ****p< 0.0001

### Effects of stress on EV production

We encountered one instance where the screen phenotype appeared to be the complete opposite of the phenotype we observed when vesicles were isolated using the conventional EV isolation protocol. The *sarA* mutant was classified as a hypervesiculator in the screen (Table 1). However, we initially found the *sarA* mutant strain to be severely hypovesiculating compared to the wild type strain in our large-scale verification (Figure 2B, shaking). SarA is a member of the SarA family of transcriptional regulators and is a global regulator of virulence factors (Cheung et al., 2008). The *sar* locus contains three promoters, P1, P2, and P3, each capable of producing a transcript encoding SarA. While the transcription of *sarA* from the P1 and P2 promoters is dependent upon the housekeeping sigma factor, sigma-70 (σ^70^), transcription from the P3 promoter requires the alternative sigma factor, σ^B^, which is activated under various stress conditions (Bischoff et al., 2001). It has been shown that the σ^B^-dependent P3 transcript is maximally expressed during late exponential phase when σ^B^ is most active. Additionally, it has also been shown that σ^B^ is required for stress resistance and the acid-adaptive response in exponential-phase cells (Chan et al., 1998). Because the timepoint we chose to assess vesicle production coincides with the maximal expression of the σ^B^-dependent *sarA* transcript, as well as the timepoint in which σ^B^ is most active, we wondered if the growth conditions of the screen had an effect on EV biogenesis. Due to the high surface area to volume ratio, limited media volume, and limited shaking/aeration of the culture in a 96-well microtiter plate, we speculated that growth in a 96-well microtiter plate may be creating an environment that is amplifying the stress experienced by the cells than when grown in a large flask. To test this, we simulated growth in a 96-well microtiter plate in a larger scale by growing the *sarA* mutant in a beaker without any shaking. Compared to the wild type strain grown in a flask and with shaking, the *sarA* mutant grown in the microtiter plate-mimicking condition exhibited a hypervesiculating phenotype just as we observed in the screen (Figure 2B, static). This result demonstrates that cellular stress promotes EV biogenesis. It is also consistent with prior studies demonstrating that σ^B^ activation and upregulation of the σ^B^-dependent stress response promote EV biogenesis (Li et al., 2024; Mitsuwan et al., 2019; Qiao et al., 2022; van Schaik & Abee, 2005). Additionally, this result highlights a shortcoming in our screen. The growth conditions in the 96-well plate-based screen we developed seem to be creating a more stressful environment than typical flask culturing conditions. This suggests that mutations that render the cells unable to handle certain stresses may have EV phenotypes that were miscategorized or excluded in our screen analyses.

Vancomycin is a glycopeptide antibiotic derived from the Gram-positive organism *Amycolatopsis orientalis* and is commonly used as a last resort antibiotic for treating *S. aureus* infections. To investigate the potential role of EVs in relieving cells from environmental stressors such as sublethal amounts of antibiotics, we grew *S. aureus* JE2 in a subinhibitory concentration of vancomycin and in the presence of varying concentrations of purified EVs isolated from JE2 overnight cultures grown in TSB. We used Gompertz least squares regression to model the growth, lag time, and maximum culture density. Treatment with the subinhibitory concentration of vancomycin alone increased the lag time of *S. aureus* growth, and decreased the maximum density achieved by the culture, as expected (Figures 3A and 3B). Interestingly, however, we noted a dose-dependent increase in the maximum culture density when EVs were added in addition to vancomycin, with the culture supplemented with 100 µg/ml EVs growing to a higher cell density compared to the no antibiotic control (Figures 3A and 3C). This suggests that, not only does EV production increase, but the presence of EVs can positively affect *S. aureus* survival in conditions of environmental stress.

**Figure 3:**
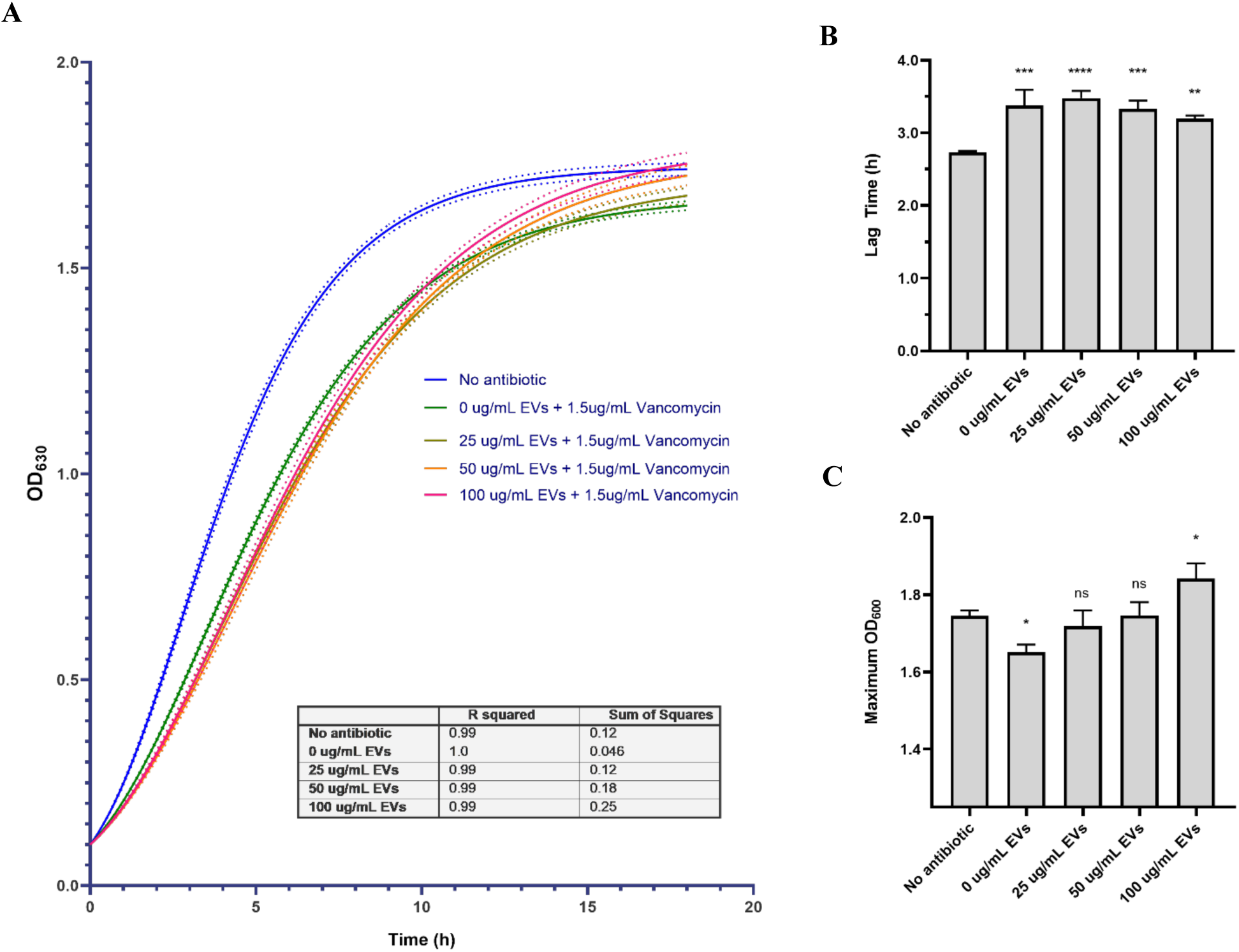
EVs increase tolerance to Vancomycin. **A.)** Best fit curves of *S. aureus* JE2 growth in TSB in the presence of 1.5µg/ml vancomycin and increasing purified EV concentrations determined by a Gompertz least squares regression model. Dashed lines represent the 95% confidence bands for the best fit curve. R^2^ values and sum of squares were determined as measured for goodness of fit. Data shown are the mean of three biological replicates. Lag times B.) and Maximum cell density values C.) were derived from best fit values determined by a Gompertz least squares regression model from three biological replicates. Values are presented as means from three independent experiments ± SD. Samples were compared using one-way ANOVA with Dunnett’s multiple comparison test. *p< 0.05, **p< 0.01, ***p< 0.001, ****p< 0.0001

### Relationship between membrane fluidity and EV production

Previously, it was shown that an increase in membrane fluidity could promote vesicle production (Schlatterer et al., 2018; Wang et al., 2020). One of the ways *S. aureus* alters its membrane fluidity is by changing the length and structure of the fatty acids incorporated into their phospholipids (Parsons & Rock, 2013; Zhang et al., 2008). Many Gram-positive bacteria, including *S. aureus*, do not produce any unsaturated fatty acids. Instead, they synthesize BCFAs from BCAAs to incorporate into phospholipids to increase membrane fluidity. Because BCFAs are derived from BCAAs, and CodY is an inhibitor for BCAA biosynthesis and import genes, we wondered if the *codY* mutant had increased membrane fluidity. Additionally, we also assessed the membrane fluidity of the other mutants to determine a possible relationship between EV biogenesis and membrane fluidity. We used the fluorescent lipophilic pyrene probe, pyrene decanoic acid (PDA), to quantitatively measure membrane fluidity. In all cases we tested, a hypovesiculating phenotype is accompanied by a decrease in membrane fluidity, and a hypervesiculating phenotype by an increase in fluidity compared to the wild type strain, suggesting that membrane fluidity is influenced by CodY, and that EV biogenesis may be regulated by membrane fluidity (Figure 2C).

### Nutritional stress promotes EV biogenesis

We observed that the *codY* mutant was hypervesiculating (Figure 2A, TABLE1). Due to its role as a repressor of genes involved in the response to BCAA limitation, the *codY* mutant strain would be overexpressing those genes as if the cell is experiencing nutritional stress. This result suggests that nutritional stress may promote EV biogenesis.

The regulation of CodY-DNA binding is dependent on BCAA availability as well as intracellular GTP levels, which is directly influenced by the stringent response. RelA/SpoT homolog (RSH) is the bifunctional stringent response regulator which synthesizes and hydrolyzes the stringent response alarmone, (p)ppGpp, directly altering the intracellular GTP pool. RSH regulates the cell’s response to amino acid starvation, specifically the branched chain amino acids leucine and valine, through CodY (Geiger et al., 2010; Geiger & Wolz, 2014). During starvation, RSH synthesizes (p)ppGpp causing a drop in cellular GTP levels. The drop in GTP level and limited BCAA availability releases CodY from binding its cognate DNA so that the CodY regulon, which includes *agr* and BCAA synthesis and transport genes, can be expressed. The RSH hydrolase activity is essential, and disruption of the hydrolase is lethal except in a (p)ppGpp null strain. The (p)ppGpp synthetase activity of RSH is restricted and is activated only upon interaction of the C-terminal domain (CTD) with ribosomal partners (Gratani et al., 2018). The *rsh* mutant from the NTML has a transposon insertion in the CTD, and it has been previously demonstrated that RSH lacking the CTD is tightly held in the hydrolase-ON state. Thus, the *rsh* mutant (which we will now refer to as the *rsh_syn_* mutant) from the NTML retains its hydrolase activity but is unable to synthesize the alarmone to activate the stringent response. We observed that the *rsh_syn_* mutant was hypervesiculating, suggesting that the inability to respond to stress through the stringent response promotes vesicle production (Figure 2A).

### Membrane components influencing EV biogenesis

The *S. aureus lgt* gene encodes a phosphatidylglycerol-prolipoprotein diacylglyceryl transferase that catalyzes the first step of lipoprotein biosynthesis (Sankaran & Wu, 1994). The *lgt* mutant is deficient in the lipidation and maturation of lipoproteins, but the proteins themselves are still produced, however, they are not retained in the bacterial membrane (Stoll et al., 2005). The *lgt* mutant has been previously shown to have a vesiculation phenotype (Wang et al., 2020). Interestingly, the prior study found the *lgt* mutant to be hypervesiculating with increased membrane fluidity compared to the wild type. However, here we found the *lgt* mutant is hypovesiculating with decreased membrane fluidity (Figures 2A and 2C). It is likely that the discrepancy in results is due to differences in growth conditions, as we had previously noted how changes in growth conditions drastically altered EV production (Figure 2B). Nevertheless, the observation of a vesiculation phenotype in the *lgt* mutant in two separate studies would suggest that the absence of lipoproteins affect vesiculation.

Wall teichoic acids (WTA) have also previously been implicated in EV biogenesis in *S. aureus*, where cells deficient in WTAs produced more EVs than the wild type strain (Wang et al., 2018). WTAs are cell wall associated glycopolymers and are a major constituent of the Gram-positive cell envelope (Brown et al., 2013). They have been shown to play a role in cell division, biofilm formation, as well as in staphylococcal adhesion and colonization. *S. aureus* makes two distinct WTA polymers, the longer TarL directed polymer (L-WTA) and the shorter TarK directed polymer (K-WTA) (Meredith et al., 2008). While the absence of TarL is lethal, TarK is dispensable and is repressed by *agr*. Agr expression is maximal during late exponential phase, thus the proportion of L-WTA to K-WTA increases as the cells reach the post-exponential phase (Podkowik et al., 2024). Previously, it was shown that the expression of the operon that encodes TarK may also be under direct regulation by CodY (Gao et al., 2023). Here we found that the *tarK* mutant, which only makes the long L-WTA polymer directed by TarL, had reduced vesicle production (Figure 2A). Therefore, it is likely the WTA chain length may influence EV production. Additionally, the temporal expression of *tarK* and the vesiculation phenotype of the *tarK* mutant suggests that vesicle production is more robust during exponential growth compared to the post-exponential phase. Finally, the dynamic regulation of *tarK* by *agr* and CodY emphasizes the role of the *agr* quorum sensing system in EV biogenesis, and further suggests that nutrient limitation, specifically BCAA limitation, plays a role in EV biogenesis.

### *Agr*-based regulation of EV production

The *agr* quorum sensing system appears to play a key role in EV biogenesis as the *agr* mutant is severely hypovesiculating (Figure 2D) (Im et al., 2017). Additionally, α-PSMs, a direct target of AgrA has been previously shown to impact vesicle production, and in the absence of CodY, a repressor of *agr*, the *codY* mutant is hypervesiculating (Figures 2A and 2D) (Briaud et al., 2021; Schlatterer et al., 2018; Wang et al., 2018). The *agr* system regulates the expression of many genes, including key virulence factors such as proteases and toxins, through the small RNA, RNAIII (Jenul & Horswill, 2019). RpiRC is a transcriptional regulator of the pentose phosphate pathway (PPP) and its expression is regulated by the *agr* system’s main effector molecule, RNAIII (Hallier et al., 2024; Majerczyk et al., 2010). It has been shown that RNAIII binds and stabilizes the *rpiRc* transcript to increase its translation. Accumulation of RpiRC then represses RNAIII transcription during the exponential-growth phase, but not during the post-exponential-growth phase (Zhu et al., 2011). In the absence of RpiRC, RNAIII transcription is increased, resulting in more virulence factor production and secretion, which in turn increase *S. aureus* pathogenicity (Balasubramanian et al., 2016; Gaupp et al., 2016). In our study, the absence of RpiRC increased both membrane fluidity and EV production of the cell (Figures 2A and 2C). The PPP is a primary source of the cofactor NADPH and precursors for important cellular processes such as nucleic acid, aromatic amino acid, and fatty acid biosynthesis. Therefore, the *rpiRc* mutant, which is unable to upregulate the expression of PPP genes, is likely to be experiencing some degree of nutritional stress in comparison the wild type strain. This further supports the hypothesis that nutritional stress is a driver for EV production. Additionally, the RNAIII-dependent regulation of *rpiRc* establishes a direct link between central metabolism, *agr* quorum sensing system, and EV biogenesis.

Another *agr* target that has previously been implicated in *S. aureus* EV biogenesis are the α-PSMs. α-PSMs are regarded as key secreted toxins by *S. aureus*, however while the expression of most of *S. aureus*’ secreted virulence factors are controlled by *agr* through RNAIII, α-PSM expression is controlled by AgrA independently of RNAIII (Jenul & Horswill, 2019; Queck et al., 2008). Additionally, modulation of its expression is determined by the RSH synthetase activity, which also influences CodY expression, AgrA expression, and thus α-PSM expression itself (Geiger et al., 2012; Jenul & Horswill, 2019). Using homologous recombination, we generated the α-PSM deletion mutant (JE2 Δ*psmα*) in the NTML parent strain JE2 and assessed its vesicle production and membrane fluidity compared to its parent strain (Figure 4A). Consistent with the *rsh_syn_* mutant, which has minimal α-PSM expression, the JE2 Δ*psmα* mutant has increased membrane fluidity accompanied by a hypervesiculating phenotype (Figures 4A, 4B, 2A, and 2C, Supplemental Figure 2) (Geiger et al., 2012). Complementation with a plasmid constitutively expressing α-PSMs reduced both EV production and membrane fluidity, demonstrating that α-PSM expression negatively influences EV production (Figures 4C and 4D). Its stringent response-dependent regulation also links the *agr* quorum sensing system and EV biogenesis to the response to amino acid limitation.

**Figure 4:**
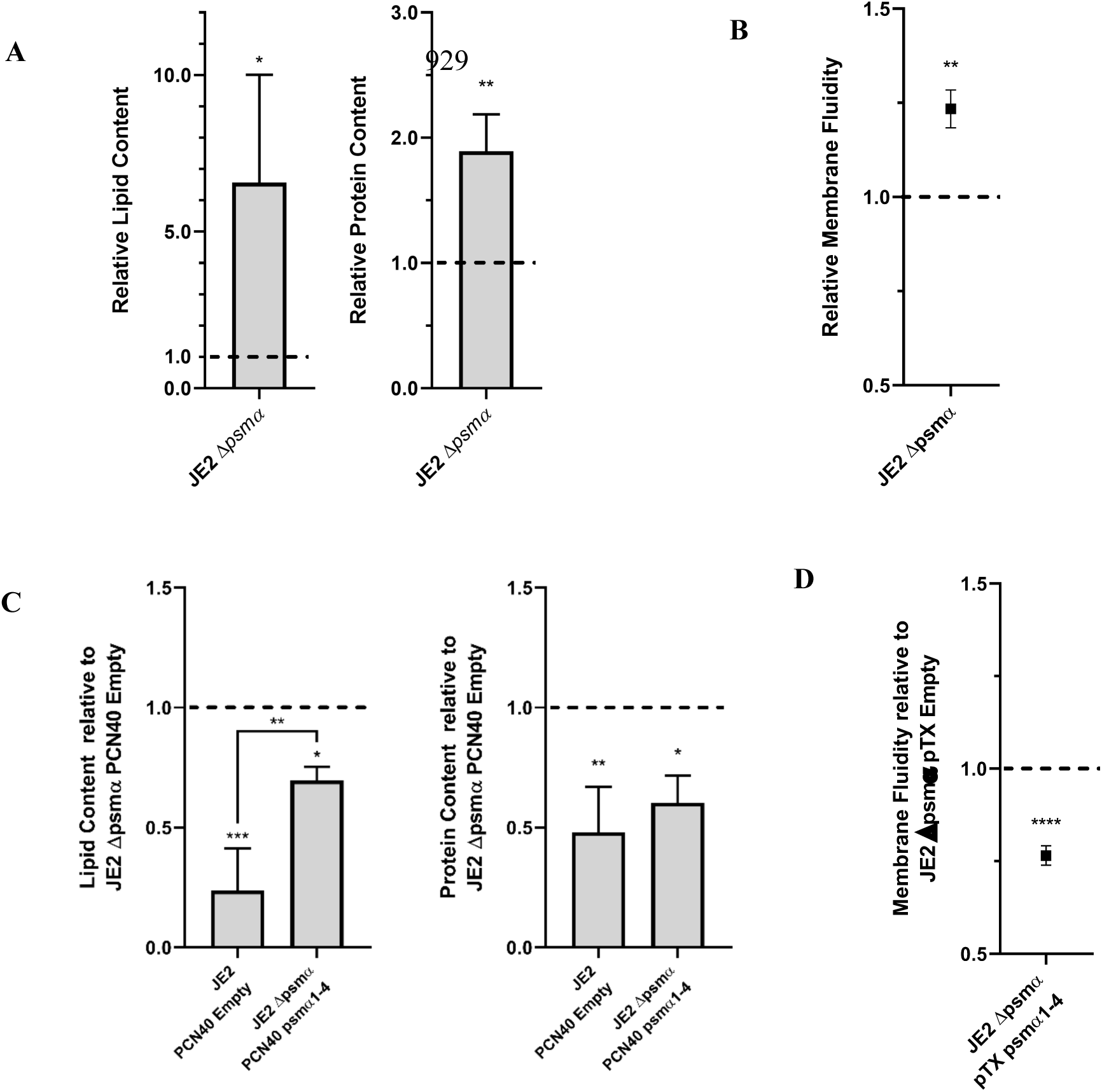
α-PSMs inhibit EV biogenesis. **A**.) The JE2 Δ*psmα* mutant is hypervesiculating compared to the JE2 wild type strain represented by the dashed lines. **B.)** The JE2 Δ*psmα* mutant has increased membrane fluidity compared to the JE2 wild type strain. **C.)** Complementation of the JE2 Δ*psmα* mutant with the PCN40 expression vector expressing α-PSMs1-4 reduces EV production compared to the JE2 Δ*psmα* mutant with the empty PCN40 expression vector represented by the dashed lines. **D.)** Complementation of the JE2 Δ*psmα* mutant with the PCN40 expression vector expressing α-PSMs1-4 has decreased membrane fluidity compared to the JE2 Δ*psmα* mutant with the empty PCN40 expression vector represented by the dashed lines. Values are presented as means from three independent experiments ± SD normalized to the control. Samples were compared using an unpaired t-test (**A**, **B**, and **D**) or using one-way ANOVA with Tukey’s multiple comparison test (**C**). *p< 0.05, **p< 0.01, ***p< 0.001, ****p< 0.0001

While the change in lipid content is typically larger than the change in protein content in the EVs of the vesiculation mutants, we observed the opposite in the *rpiRc* mutant (Figure 2A). We speculated that this is due to the increased RNAIII expression resulting in increased virulence factor production and secretion reported in the absence of RpiRC (Balasubramanian et al., 2016; Gaupp et al., 2016). We also wondered if the increased RNAIII expression is contributing to the hypervesiculating phenotype of the *rpiRc* mutant. Deletion of *rnaiii* by allelic exchange resulted in a hypovesiculating phenotype, suggesting that RNAIII expression promotes vesiculogenesis (Figure 5).

**Figure 5:**
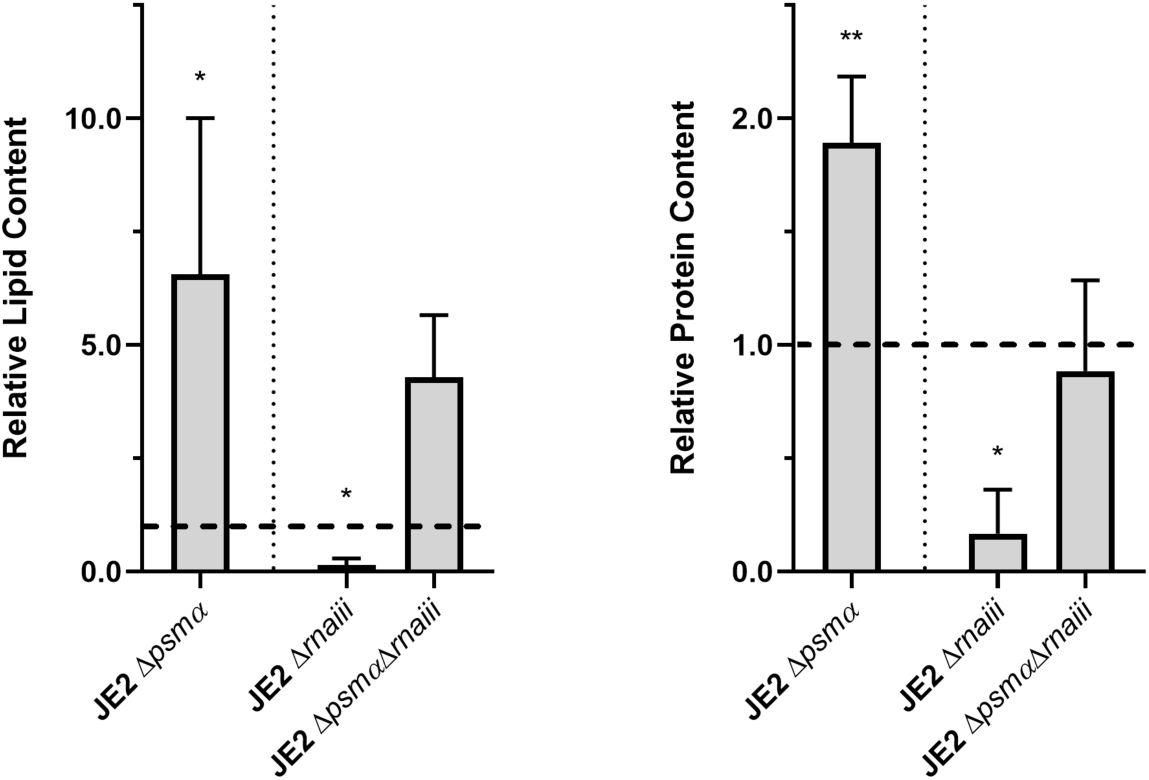
RNAIII and α-PSMs have opposing contributions to EV biogenesis. The JE2 Δ*rnaiii* mutant is hypervesiculating compared to the JE2 wild type strain, while the JE2Δ*psmα*Δ*rnaiii* mutant’s vesiculation is not significantly different from the JE2 wild type strain represented by the dashed lines. The JE2 Δ*psmα* mutant on the left is the same as in Figure 4A and was only added for comparison and was not included in the analysis. Values are presented as means from three independent experiments ± SD. Samples were compared using one-way ANOVA with Dunnett’s multiple comparison test. *p< 0.05, **p< 0.01, ***p< 0.001, ****p< 0.0001

RNAIII and α-PSMs are direct targets of AgrA, however deletion of *rnaiii* results in a hypovesiculating phenotype similar to the *agr* mutant, while the *psmα* deletion resulted in a hypervesiculating phenotype similar to the mutants experiencing nutritional stress. Intriguingly, deletion of both *rnaiii* and *psmα* resulted in an intermediate phenotype suggesting that the effects of RNAIII and α-PSMs on EV biogenesis are independent of each other; and that EV biogenesis through the *agr* quorum sensing system is facilitated through RNAIII (Figure 5).

### Effects of α-PSM expression

Since *psmα* expression is regulated by RSH and RSH activity is governed by nutrient availability, we wondered if the hypervesiculating phenotype of the Δ*psmα* mutant is caused by nutritional stress. We supplemented the JE2 Δ*psmα* mutant with the BCAAs valine and leucine (455µM), glucose (7mM), or both to determine if increasing nutrient availability can reduce EV production. As shown in Figure 6A, supplementation of the JE2 Δ*psmα* mutant with BCAAs or glucose alone does not cause a significant change in EV production compared to the JE2 Δ*psmα* mutant, but addition of both significantly decreased EV production by 15.28% ±0.04 by lipid measure and 19.48% ±0.23 by protein measure (Figure 6A). This demonstrates that nutrient availability, indeed, has an influence on EV production, and nutrient availability is inversely related to EV production. This also supports the hypothesis that EVs are produced in response to nutrient stress. Finally, this suggests that in the absence of α-PSMs, cells are experiencing or responding to nutritional stress or apparent nutritional stress, presumably through the stringent response, causing the increase in EV production.

**Figure 6:**
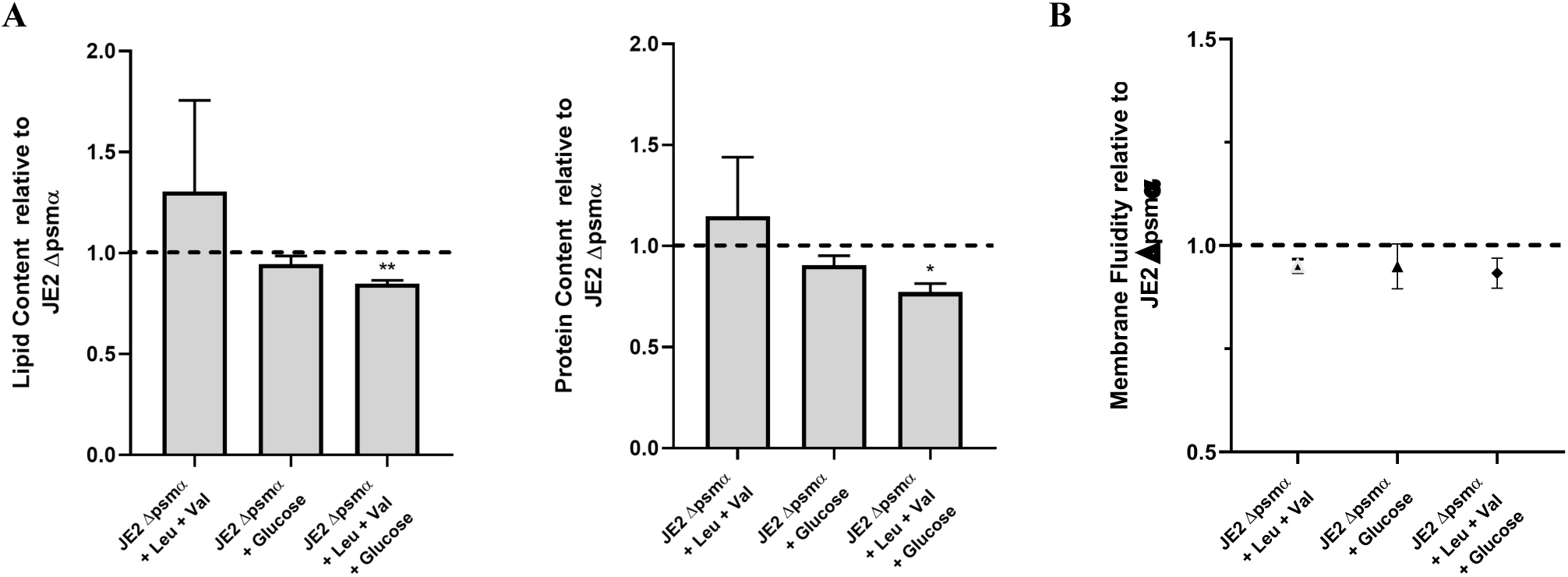
Effects of glucose and BCAA supplementation on EV production of the JE2 Δ*psmα* mutant. **A.)** Supplementation with 455µM each of leucine and valine and 7mM glucose reduces EV production in the JE2 Δ*psmα* mutant compared to the JE2 Δ*psmα* mutant without supplementation **B.)** Glucose and BCAA supplementation does not significantly reduce membrane fluidity of the JE2 Δ*psmα* mutant compared to the JE2 Δ*psmα* mutant without supplementation. Values are presented as means from three independent experiments ± SD. Samples were compared using one-way ANOVA with Dunnett’s multiple comparison test. *p< 0.05, **p< 0.01, ***p< 0.001, ****p< 0.0001

We did not observe a significant decrease in membrane fluidity despite the observed decrease in EV production when supplementing with BCAAs and glucose (Figure 6B). The cell membrane is limited to a narrow window of fluidity to remain viable and the JE2 Δ*psmα* mutant is severely hypervesiculating, and it would still be a hypovesiculating compared to the wild type even with a 15-20% decrease in its EV production (Figures 4A, 4B, 6A and 6B).Therefore, it is likely that any changes in membrane fluidity would not be drastic enough to be accurately detected by the fluorescent probe.

Previous studies suggested that α-PSMs promote EV biogenesis and increase membrane fluidity by acting on the cell membranes through their surfactant-like activity (Briaud et al., 2021; Schlatterer et al., 2018; Wang et al., 2020; X. Wang et al., 2021; Wang et al., 2018). Here, we show the opposite result: We demonstrate that α-PSMs obstruct EV biogenesis (Figure 4). In addition, the absence of α-PSMs increases membrane fluidity and changes the fatty acid composition of the membrane phospholipids shifting to more short and branched chain fatty acids (Supplemental Figure 2). We speculated that the effect of α-PSMs on EV biogenesis may be strain specific. Although the NTML parent strain JE2 is also a USA300 strain, JE2 has been cured of its plasmids, and we cannot disregard that difference (Fey et al., 2013). Thus, we acquired the α-PSM mutant and parent strain, as well as the complemented strains, used in the prior studies (Schlatterer et al., 2018; X. Wang et al., 2021; Wang et al., 2018). In our growth conditions, the USA300 LAC Δ*psmα* mutant is hypervesiculating, and complementation with α-PSMs decrease EV production, just as we observed in the JE2 Δ*psmα* mutant (Figure 4 and Supplemental Figure 3). It is possible that differences in growth conditions and/or EV isolation methods may be responsible for the differences in our results – we have previously demonstrated that growth conditions can alter EV production significantly (Figures 2B and 7). More recently, a different group described a hypervesiculating mutant in *S. aureus*. Deletion of the gene that encodes Asp23, which is involved in stress response in alkaline conditions, resulted in a hypervesiculating phenotype (Li et al., 2024). RT-qPCR analysis revealed that their hypervesiculating mutant had lowered *psmα* expression, and complementation of the Δ*asp23* mutant with α-PSMs reduced EV production, similar to the results presented in this study.

**Figure 7:**
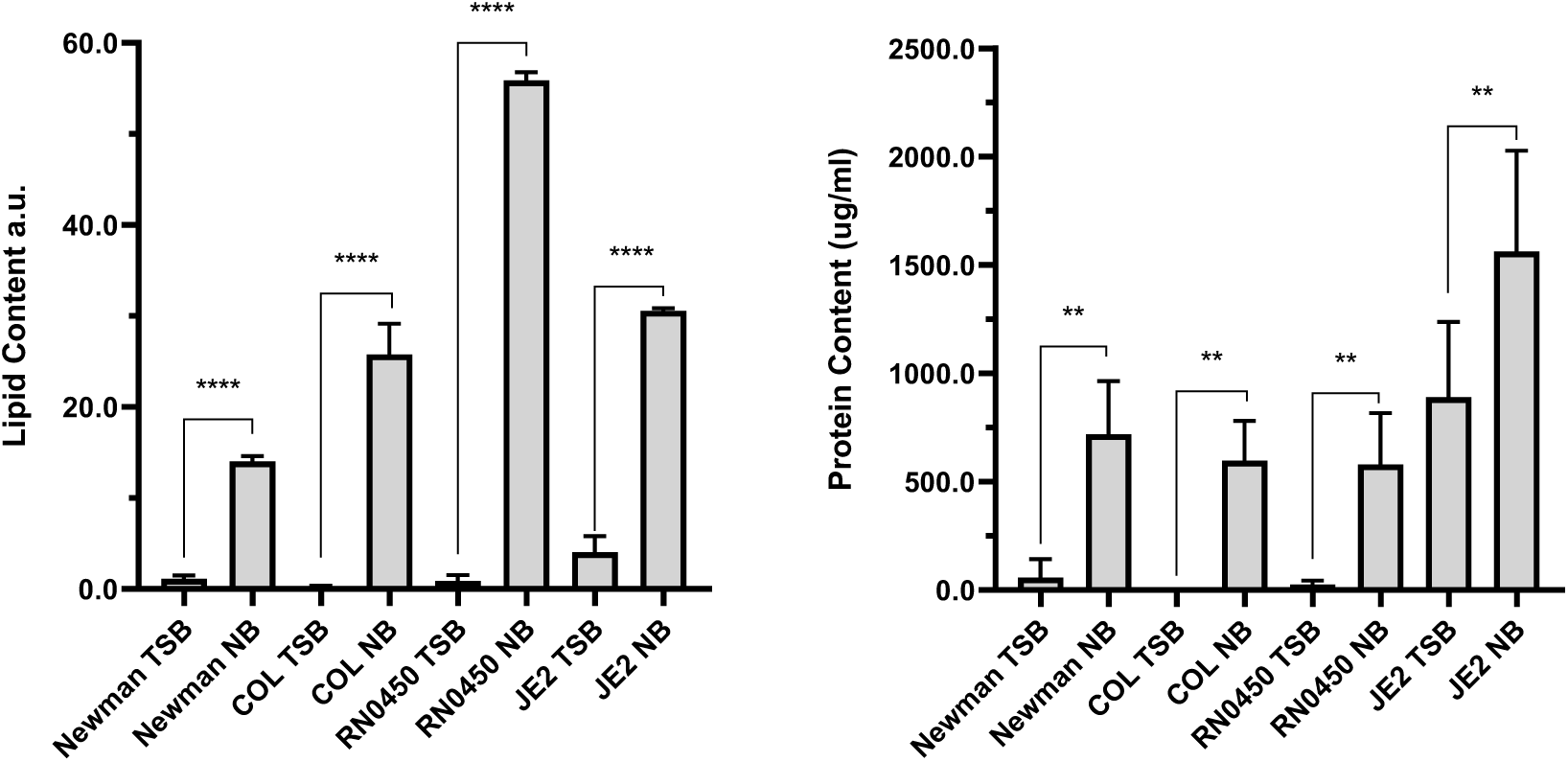
*S. aureus* vesicle production in rich or moderately rich growth medium. EVs purified from MRSA strains, COL and JE2, and MSSA strains, Newman and RN0450, grown for 18h in rich medium (TSB) or moderately rich medium (NB). Values are presented as means from at least three independent experiments ± SD. Samples were compared using one-way ANOVA with Sidak’s multiple comparison test. *p< 0.05, **p< 0.01, ***p< 0.001, ****p< 0.0001

### Nutrient availabity and vesiculogenesis

To determine if the relationship between nutrient availability and EV production is conserved between strains, and to further define the role of *agr* in this process, we grew *S. aureus* JE2, a methicillin resistant *S. aureus* (MRSA) strain, and several other *S. aureus* strains in rich media (TSB) or a moderately rich growth medium (NB) and assessed their vesicle production. Vesicles were isolated from cultures grown overnight (18h) to ensure all strains have reached stationary phase and to maximize nutrient depletion in the media. COL is a MRSA strain that has a defect in *agr*, causing low *agr* expression, and RN0450 and Newman are both methicillin sensitive *S. aureus* strains (MSSA) (Herbert et al., 2010). RN0450 has a deletion in the σ^B^ positive regulator, RbsU; and Newman carries a mutation causing the constitutive activation and upregulation of the CodY-regulated SaeRS two-component system, which controls a different set of virulence factors as *agr*. We observed that all strains produced more vesicles in the moderately rich growth medium compared to the rich growth medium, further demonstrating that limited nutrient availability promotes vesicle formation (Figure 7).

Additionally, COL, which has low *agr* expression, barely produced any vesicles when grown in rich media, but produced significantly more when nutrients are limited, but not to the extent of the other MRSA strain JE2; solidifying the role of *agr* in EV biogenesis during nutrient limitation (Figure 7). The MSSA strains Newman and RN0450 did not produce as much vesicles as the MRSA strain JE2 when grown in rich media, however when grown in the moderately rich media, RN0450 produced more vesicles, by lipid measurement, than JE2 grown in moderately rich media, suggesting that EV production, indeed is made in response to nutrient stress, but not exclusively to nutrient stress (Figure 7). Finally, the Newman strain significantly increasing vesicle production when there is less nutrient availability suggests that Sae activation is not a major contributor to EV biogenesis during nutrient limitation despite being part of the CodY regulon (Figure 7).

The *agrA* mutant was previously shown to be hypovesiculating (Im et al., 2017; Wang et al., 2018). Isolation and quantification of vesicles from the *agrA* mutant, indeed, show that it is a hypovesiculator (Figure 2D), however the AgrA mutant was not identified in our screen to have a vesiculation phenotype. This result drew our attention to the impact of our screen design on the evaluation of MAD values. First, the phenotypic assessment of mutants within a single plate could be biased in one direction or the another. Specifically, the MAD values were calculated from the values within each specific plate in order to minimize errors that would arise from day-to-day differences in the fluorescence intensities of the various fluorescent probes used in the screen. However, this was based on the key assumption that the average vesiculation value of each plate would be representative of wild type vesiculation. Therefore, if a particular plate is skewed to contain more mutants of one vesiculation phenotype, this could exclude some mutants with the opposite phenotype from being identified in the screen. Second, the selected vesiculation cutoff of 2-MADs away from the mean normalized vesiculation value was arbitrarily assigned during the design of the screen. For the *agrA* mutant to be considered a hypovesiculator in our screen, the MAD cutoff would have to be adjusted to 1.2-MADs away from the mean vesiculation value. When we adjusted the stringency to 1.5 and 1.2-MADs away from the mean vesiculation value, we identified an additional 207 and 435 (207 plus 228) mutants, respectively, to have vesiculation phenotypes (Supplemental Tables 3 to 6).

We decided to isolate and assess EV production from several mutants identified using these less stringent parameters to determine if these less stringent cutoff parameters can still successfully identify mutants with vesiculation phenotypes. While the less stringent parameters successfully identified hypervesiculators, it was less reliable with hypovesiculators. Three out of the three selected hypervesiculators were also hypervesiculators when we assessed EVs isolated using the conventional ultracentrifugation-based EV isolation protocol, while only one out of the three selected hypovesiculators had a phenotype consistent with the screen (Figure 8A).

**Figure 8:**
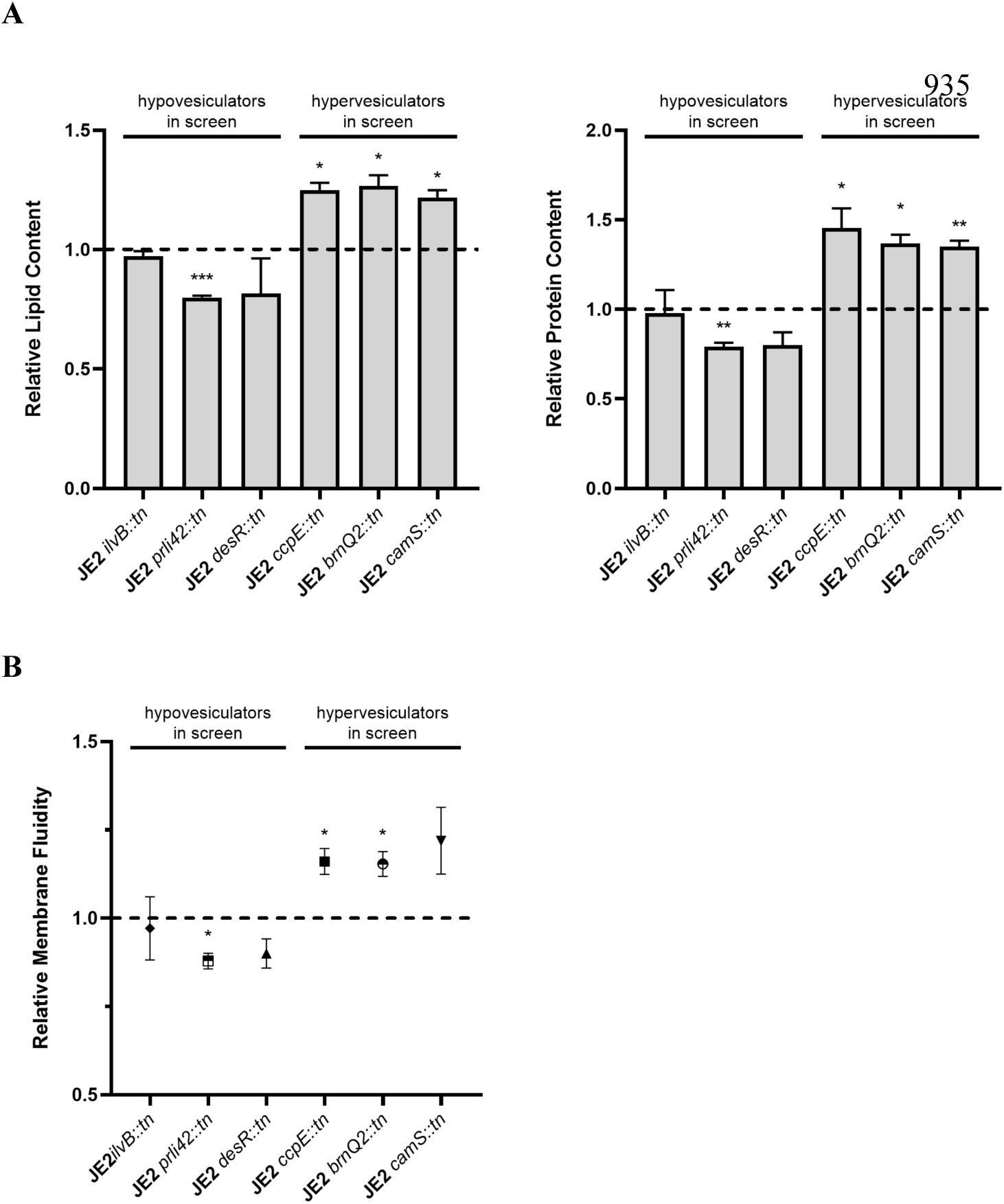
Decreasing stringency of screen parameters still successfully identify hypervesiculators. **A.)** Lipid and protein contents of EVs purified from mutants were compared to that of the wild type JE2 parent strain represented by the dashed lines. **B.)** Relative membrane fluidity of mutants were compared to that of the wild type JE2 parent strain represented by the dashed lines. Values are presented as means from three independent experiments normalized to the control ± SD. Samples were compared using mixed effects analysis with Dunnett’s multiple comparison test. *p< 0.05, **p< 0.01, ***p< 0.001, ****p< 0.0001

Membrane fluidity measurements further support that EV biogenesis is correlated with the fluidity of the membrane (Figure 8B). This suggests that the hypovesiculating mutants identified in the screen using more lenient parameters may not be as reliable as the hypervesiculators.

## Discussion

In this study, we designed a moderately high-throughput screen for vesiculation and demonstrated that it successfully identified mutants in the NTML with vesiculation phenotypes. To our knowledge, this is the first genome-wide screen to identify genetic determinants of EV production in *S. aureus*. In analyzing and validating the screen results, we learned that it is important to verify EV phenotypes using an alternate culturing and assessment method, as the growth conditions in the screen (e.g. gas exchange, limited medium, limited shaking/nutrient distribution) may impact EV production. In addition, because the screen relies on a number of lipophilic fluorescent probes, it is likely that some mutations that cause changes in the properties of the cell membrane can also change how the dyes interact them. Nevertheless, the results provide validated, and important insight into *S. aureus* genes and pathways affecting EV production.

The results of our screen implicate 173 genes influencing vesiculogenesis in *S. aureus*. Additionally, our data point to CodY as a mediator to EV biogenesis and suggest that nutrient limitation is a driver for vesicle production (Figure 9). We showed that the CodY-regulated *agr* quorum sensing system is a key mediator to EV production and that the AgrA targets, α-PSMs and RNAIII, have opposing contributions to EV biogenesis. The vesiculation phenotypes of strains with transposon insertions in the genes that encode the *αpsm and rnaiii* regulators, RSH and RpiRC, establishes, for the first time, a link between the stringent response and PPP to the quorum sensing system in regulating vesiculogenesis through α-PSMs and RNAIII. Overall, our results demonstrate that vesicle production is a response to cellular stress, including nutritional stress, and is largely regulated and orchestrated by the *agr* quorum sensing system, possibly through crosstalk with other cellular pathways such as the stringent response through α-PSMs, and central metabolism (i.e. PPP) through RNAIII.

**Figure 9:**
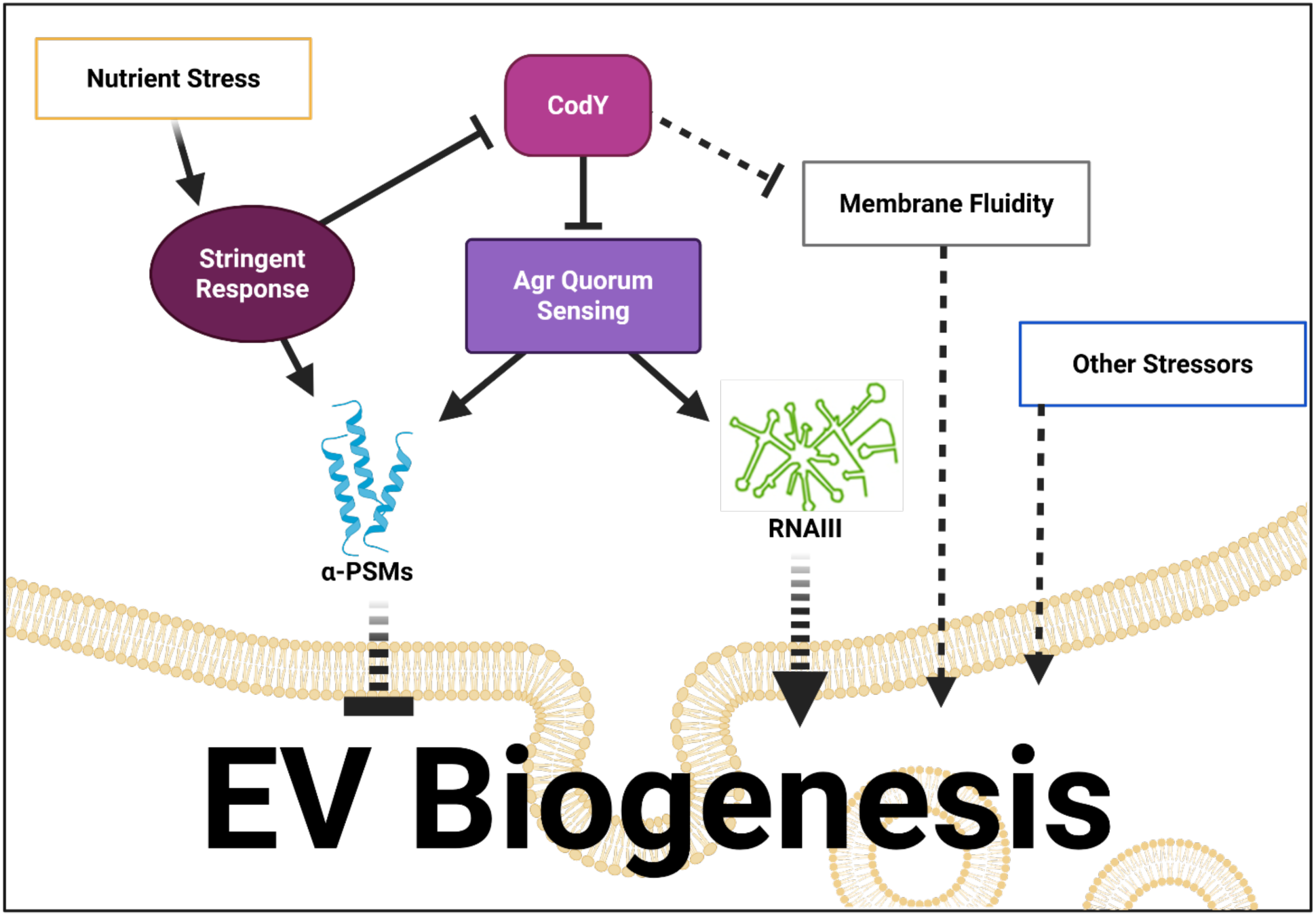
Model for regulation of EV biogenesis in *S. aureus.* Nutrient limitation promotes EV production presumably by activating the stringent response, which promotes the expression of CodY-regulated genes and activation of the *agr* quorum sensing system. The *agr* system largely controls EV biogenesis through RNAIII. In addition to AgrA, the stringent response regulator, RSH, regulates the expression of α-PSMs, whose effect on EV biogenesis is opposite and independent of RNAIII. Additionally, membrane fluidity, which can be influenced by CodY through BCAAs is correlated with EV biogenesis.

Production of vesicles is a universally conserved process, suggesting the existence of a conserved mechanism or principle for EV biogenesis. The stringent response is a conserved bacterial stress response and recent advances in stringent response research have shown a recurring theme. Several studies have shown ribosome-independent stimulation of the stringent response by direct binding of small regulatory proteins to RSH to control cellular (p)ppGpp levels. In *E. coli*, the accumulation of (p)ppGpp during fatty acid starvation required the interaction between the acyl carrier protein and the RSH enzyme, SpoT (Battesti & Bouveret, 2006). Similarly, a small protein expressed during phosphate starvation, YtfK, was shown to promote the synthesis of (p)ppGpp during phosphate and fatty acid starvation by direct interaction with SpoT (Germain et al., 2019). Rsd, a protein originally identified as an antagonist of the housekeeping sigma factor, σ^70^, was shown to interact with SpoT and activates its (p)ppGpp hydrolase activity in response to a nutrient downshift from a preferred carbon source to a less preferred one (Lee et al., 2018). More recently, NirD, the small subunit of the nitrite reductase, was demonstrated to inhibit the (p)ppGpp synthetase activity of the RSH enzyme, RelA, through direct interaction with the catalytic domains of RelA (Leger et al., 2021). In C. *crescentus*, the phosphorylated EIIA^Ntr^ enzyme of the nitrogen-sensing phosphotransferase system directly binds the ACT domain of SpoT to inhibit the hydrolysis of (p)ppGpp during nitrogen starvation (Ronneau et al., 2019). In *V. cholerae*, depletion of the protein CgtA caused changes in gene expression consistent with induction of the stringent response, and a two-hybrid assay showed that CgtA interacts with SpoT (Raskin et al., 2007). In the Gram-positive bacterium *B. subtilis*, a protein with previously unknown function, DarB, was demonstrated to directly interact with Rel during potassium limitation and low intracellular levels of cyclic di-AMP to promote the accumulation of (p)ppGpp (Ainelo et al., 2023; Kruger et al., 2021).

In *S. aureus*, most genes that are activated during the stringent response are regulated through CodY derepression (Geiger et al., 2012). Geiger and colleagues identified 150 (p)ppGpp dependent genes whose upregulation during the stringent response was dependent on CodY. In contrast, they found only 11 (p)ppGpp dependent genes that were upregulated during the stringent response independently of CodY, four coded for the α-PSMs and two for the β-phenol soluble modulins (β-PSM1, and β-PSM2, collectively β-PSMs). Similarly, majority of the genes activated by the *agr* quorum sensing system are regulated through RNAIII, while genes upregulated by the *agr* independently of RNAIII almost exclusively comprised the *psmα* and *psmβ* genes (Queck et al., 2008). Interestingly, *S. aureus* possesses another α-PSM, delta hemolysin (*hld*), whose expression is regulated by RNAIII (Cheung et al., 2014). And while the *psmα* locus is conserved in all *S. aureus*, including non-pathogenic strains; *hld* is present mostly in pathogenic strains.

Given the hypervesiculating phenotype of the Δ*psmα* and *rsh_syn_* mutants, the RNAIII-independent and RSH-dependent regulation of α-PSMs, the conservation of the *psmα* locus, the additive effect of *psmα* and *rnaiii* deletion on EV biogenesis, and the ability to decrease vesicle production of the Δ*psmα* mutant by nutrient supplementation, it is possible that α-PSMs may be regulators of the stringent response (Figures 2A, 4A, and 5). Based on our results, it is possible that α-PSMs act as a regulatory feedback inhibitor to RSH to fine tune the stringent response, and α-PSMs can regulate the degree of EV production by *agr* based on nutrient availability through the stringent response. This is interesting and important to address in future studies and may uncover a conserved principle of EV biogenesis in bacteria.

Many of the RNAIII-dependent genes upregulated by *agr* are secreted virulence factors and have been found packaged in vesicles, and deletion of *rnaiii* reduces vesiculation to almost undetectable levels, similar to the *agr* mutant (Figures 5, 2A, and 2D) (Askarian et al., 2018; Gurung et al., 2011; Im et al., 2017; Lee et al., 2009; Schrempf et al., 2011; Wang et al., 2018). Additionally, the *S. aureus* COL strain, which has been previously been shown to have low RNAIII levels, produce very little vesicles when grown in rich medium, and lower EV production compared to another MRSA strain, JE2, when grown in moderately rich medium (Figure 7) (Herbert et al., 2010). Thus, it seems that EV biogenesis directed by the *agr* quorum sensing system occurs through RNAIII. RNAIII has also been shown to regulate the translation of the PPP regulator *rpiRc*, while RpiRC accumulation downregulates *rnaiii* transcription (Hallier et al., 2024; Majerczyk et al., 2010; Zhu et al., 2011). Moreover, in the absence of RpiRC, RNAIII transcription is increased, resulting in an increase in virulence factor production and secretion, and thus pathogenicity (Balasubramanian et al., 2016; Gaupp et al., 2016). The PPP is crucial for metabolic flexibility and is a major source of NADPH required for lipogenesis, and precursors for important cellular processes such as nucleic acid, and aromatic amino acid biosynthesis. Thus, it is possible that RNAIII can also regulate the degree of EV production based on the cell’s metabolic status. This is an interesting idea and warrants further investigation.

## Materials and Methods

### Bacterial strains and plasmids

*S. aureus* strains were grown in Tryptic Soy Broth (TSB, Millipore 22092) at 37°C with shaking at 250 rpm unless stated otherwise. *E. coli* strains were grown in Luria-Bertani (LB, Millipore VM782285) broth at 37°C with shaking at 250 rpm. Antibiotics were added in the following concentrations when appropriate: *S. aureus*: Chloramphenicol (Fisher Scientific, BP904, 10µg/ml), Erythromycin (Sigma, E5389, 10µg/ml), Anhydrotetracycline (Thermo Scientific, J66688.MB, 1µg/ml), Tetracycline (Sigma, T3258, 10µg/ml) *E. coli*: Chloramphenicol (Fisher Scientific, BP904, 10µg/ml), Ampicillin (Sigma, A6140, 100µg/ml). All bacterial strains and plasmids used in this study are listed in Table 4.

**Table 4:**
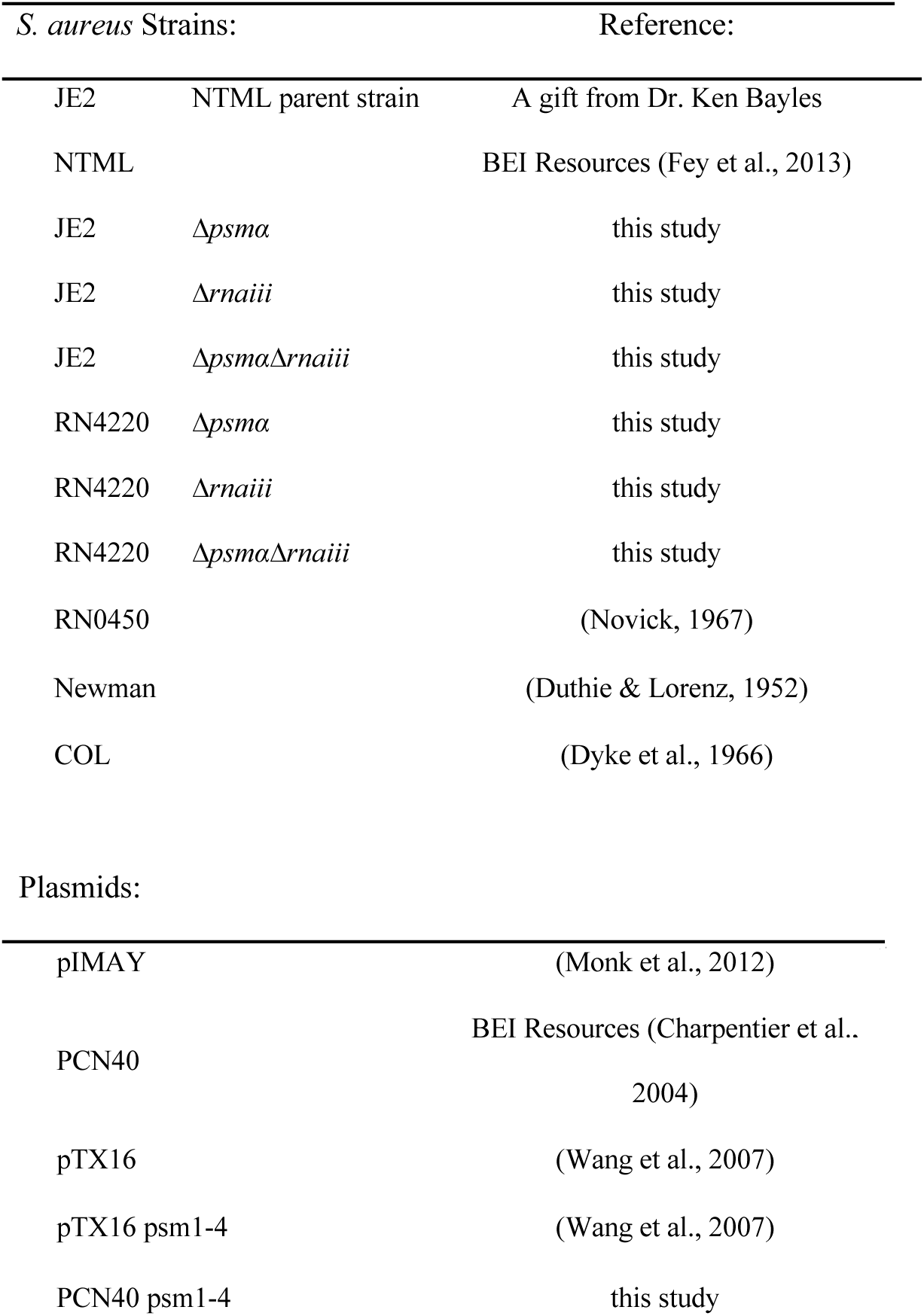
Strains and plasmids used in this study.

### Screen development

To compare the LIVE/DEAD BacLight (Invitrogen L13152) assay with SYBR Gold/PI viability assay, *S. aureus* JE2 was grown for 5 hours to early stationary/late exponential phase. Half of the culture was heat killed by boiling for 20 mins and assigned to be a 0% live culture. The other half of the culture was assigned to be the 100% live culture. Both cultures were mixed in various ratios and diluted 2-fold or 10-fold with TSB. For SYBR Gold/PI: 10 µl SYBR Gold (Invitrogen S11494; 10,000 × stock) was mixed with 60 µl propidium iodide (PI; 20 mM stock, Invitrogen) into 1.0 ml of sterile TE buffer and vortexed thoroughly.10 µl of the staining mixture was added to each well and mixed thoroughly. The plate was incubated at 37°C in the dark for 15 minutes. Fluorescence was monitored by excitation at 488nm and taking emission values at 635nm for propidium iodide (PI) and 535nm for SYBR Gold (Molecular Devices SpectraMax). The LIVE/DEAD assay was performed per the manufacturer’s protocol (Invitrogen L13152). Fluorescence was monitored by excitation at 488nm and taking emission values at 635nm for PI (DEAD) and 500nm for SYTO9 (LIVE) (Molecular Devices SpectraMax). To determine FM4-64 sensitivity to detect EVs, purified *S. aureus* JE2 EVs were diluted with TSB to varying concentrations as determined by protein content quantified by Bradford Assay (Bio-Rad). 1μg/ml, 2.5μg/ml, and 5μg/ml concentrations of FM4-64 (Invitrogen, T13320) were used to detect lipid content of EVs. FM4-64 fluorescence was measured by excitation at 506nm, and reading emission values at 750nm (Molecular Devices SpectraMax). JE2 growth was assessed by dilution of JE2 overnight culture 1:40 with pre-warmed TSB into a 96-well flat bottom microtiter plate. OD_630_ was monitored in a BioTek ELx808 microplate reader (Agilent) with continuous shaking at 37°C. First two rows and columns of the 96-well microtiter plate were filled with water to minimize evaporation, and each biological replicate was performed with at least three technical replicates.

### Moderately high throughput vesiculation screen

Overnight NTML mutant library strain cultures were diluted 1:40 with pre-warmed media and were grown for 5 hours in 96-well microtiter plates enclosed in a humidity chamber at 37°C with shaking (250 rpm). A 20 µl aliquot of each of the cultures were diluted 10x in TE buffer (10mM Tris-HCl, 1mM EDTA, pH 8) in a black clear bottom 96 well microtiter plate before OD_600_ was measured. Cell viability was measured from the 10x diluted cells as described previously (Feng et al., 2014) replacing SYBR Green I with SYBR Gold (Invitrogen, S11494). Briefly, SYBR Gold and Propidium Iodide (PI, Invitrogen, P1304MP) were used for double staining of nucleic acids. 10 µl SYBR Gold (10,000 × stock) was mixed with 60 µl PI (20 mM stock) into 1.0 ml of sterile TE buffer and vortexed thoroughly. 10 µl of the staining mixture was added to each well and mixed thoroughly. The plate was incubated at 37°C in the dark for 15 minutes. Fluorescence was monitored by excitation at 488nm and taking emission values at 535nm (SYBR Gold) and 635nm (PI) (Molecular Devices SpectraMax). Viability was calculated as a ratio of SYBR/Gold/PI fluorescence. The remaining cultures were sterile-filtered through 96-well, 0.45μm, PVDF, filter plates (Millipore, MSHVS4510) into 96 well black clear bottom microtiter plates. Vesiculation was measured by incubation of the filtered culture media containing secreted extracellular materials, including EVs, with 2.5µg/ml FM4-64 (Invitrogen, T13320) in the dark at 37°C for 30 mins. FM4-64 fluorescence was measured in a black clear bottom 96-well microtiter plate by excitation at 506nm, and reading emission values at 750nm (Molecular Devices SpectraMax). FM4-64 fluorescence was then normalized to OD_600_ of culture at time of harvest (Molecular Devices SpectraMax). The parameters used to define significant hypo- or hyper-vesiculation phenotypes are as follows: 1.) Cell viability value (SYBR Gold/PI) is more than 2-mean average deviations (MAD) below the mean viability value of the strains within the plate 2.) Vesiculation value (FM4-64) normalized to growth (OD_600_) is over 2-MADs from the mean normalized vesiculation value (FM4-64/OD_600_) of the strains within the plate. MAD was calculated using the formula below where x_i_ is the normalized vesiculation value of each mutant, µ is the mean of all the normalized vesiculation values, and N is the total number of mutants.

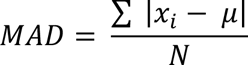

### Isolation of EVs

Overnight *S. aureus* cultures were diluted to OD_600_ = 0.1 with pre-warmed media, and grown for 5 hours. OD_600_ of cultures were measured prior to harvesting by centrifugation at 10,000 rpm (Beckman Avanti J25, JLA-10.500 rotor) at 4°C and filtration through a 0.22µm polyvinylidene fluoride (PVDF) filter (Merck Millipore Germany). Filtrates were then centrifuged at 200,000×g (Beckman Optima LE80K Ultracentrifuge, Type 50.2 Ti fixed-angle rotor) for 3 hours at 4°C. Resulting vesicle pellets were gently rinsed before being resuspended in HEPES-NaCl (10mM HEPES, 150mM NaCl, pH 7) buffer and quantified.

### Isolation of EVs for FM4-64 sensitivity assay, vancomycin tolerance assay and EV yield in TSB and NB comparison

EV isolation was performed as previously described (Lee et al., 2009). Overnight *S. aureus* cultures grown in TSB or Nutrient Broth (NB; 0.5% peptone, 0.3% beef extract, pH 7) were diluted 1:1000 in TSB or NB. Cultures were grown for 18h, at 37°C, with shaking (250 rpm). Cultures were then centrifuged at 10,000 rpm (Beckman Avanti J25, JLA-10.500 rotor) at 4°C. The cell-free supernatant was filtered through a 0.45 µm pore size, hydrophilic PVDF (Merck Millipore Germany). Filtered supernatants were concentrated by ultrafiltration using a tangential flow system (Cole-Parmer MasterFlex) with 100-kDa hollow-fiber membrane (Pall Corporation) followed by filtration through a 0.22µm PVDF filter (Merck Millipore Germany). Filtrates were then subjected to ultracentrifugation at 200,000×g for 3 h at 4°C (Beckman Optima LE80K Ultracentrifuge, Type 50.2 Ti fixed-angle rotor). Pelleted vesicles were resuspended in a small volume of HEPES-NaCl buffer before being subjected to density gradient centrifugation.

### Density gradient centrifugation

Isolated EVs were adjusted to 50% (vol/vol) Optiprep (Sigma) solution and layered at the bottom of 13.2 ml ultracentrifuge tubes. Samples were then overlaid with a 40% Optiprep layer and a 10% Optiprep layer. The tubes were ultracentrifuged at 130,000×g (Beckman Optima LE80K Ultracentrifuge, SW 41Ti Swinging-bucket rotor) for 16–18 h at 4 °C and 2-ml fractions were collected from the top of the density gradient. The Optiprep solution was removed by washing each fraction with HEPES-NaCl buffer and ultracentrifugation at 200,000×g for 3 h at 4°C (Beckman Optima LE80K Ultracentrifuge, Type 50.2 Ti fixed-angle rotor). Each fraction was resuspended in HEPES-NaCl buffer and assayed for protein concentration and lipid content using a Bradford protein assay (Bio-Rad) and 5µg/mL of FM4-64 dye (Invitrogen, T13320) respectively. EV fractions containing high proteolipid content were combined, and subject to another round of ultracentrifugation (Beckman Optima TLX ultracentrifuge, TLA-100.3 rotor; 200,000 x g, 1 h, 4°C). Resulting vesicle pellets were resuspended in an equal volume (1mL) of HEPES-NaCl buffer before being quantified.

### Quantification of EVs

For the quantification of lipid content, EVs were incubated with 5µg/mL of FM4-64 (Invitrogen, T13320) in the dark at 37°C for 30 mins. FM4-64 fluorescence was measured in a black clear bottom 96-well microtiter plate by excitation at 506nm, and reading emission values at 750nm (Molecular Devices SpectraMax). FM4-64 fluorescence was then normalized to OD_600_ of culture at time of harvest. Protein content of the vesicles were quantified by a Bradford Assay (Bio-Rad).

### Vancomycin tolerance assay

*S. aureus* JE2 overnight culture was diluted 1:40 with pre-warmed TSB supplemented with 1.5µg/ml vancomycin (Sigma, 94747) and increasing total protein concentrations of *S. aureus* JE2 EVs isolated from 18h cultures grown in TSB. Cultures were grown in a 96-well microtiter plate and growth was monitored by OD_630_ in a BioTek ELx808 microplate reader (Agilent) for 18h with continuous shaking at 37°C. First two rows and columns of the 96-well microtiter plate were filled with water to minimize evaporation, and each biological replicate was performed with at least three technical replicates.

### Membrane fluidity

*S. aureus* cultures were normalized to OD_600_=1. Cells from 1mL of OD_600_=1 cultures were resuspended in 5 µg/ml of pyrenedecanoic acid (PDA, Abcam, ab274309) and 0.08% pluronic F-127 (Sigma, P2443) in phosphate buffered saline (PBS) and stained in the dark for 30 mins at room temperature with gentle rocking. Cells were rinsed with PBS twice and resuspended in PBS before transferring to a black clear bottom 96-well microtiter plate. PDA fluorescence was monitored by excitation at 360nm and taking emission values at 400nm (monomer) and 470nm (excimer) (Molecular Devices SpectraMax). Relative membrane fluidity was calculated as a ratio of excimer to monomer fluorescence.

### DNA manipulation

We generated the JE2 *Δpsmα*, JE2 *Δrnaii*, and JE2 *ΔpsmαΔrnaiii* strains by allelic exchange as described previously (Monk et al., 2012). *S. aureus* RN4220 was used as a shuttle strain before transformation of the pIMAY plasmid into JE2 All deletion strains were confirmed by analytical PCR with genomic DNA. α-PSM complementation was done using the α-PSM-overexpressing plasmid pTx_Δ16_ psmα (Wang et al., 2007) or by PCN40 psmα generated here using the PCN40 expression plasmid constructed by Charpentier and colleagues (Charpentier et al., 2004). All *S. aureus* transformation was performed by electroporation. Preparation of electrocompetent *S. aureus* and electroporation were performed as described by Schenk and Ladagga (Schenk & Laddaga, 1992). All primers used in this study is detailed in Table 5.

**Table 5:**
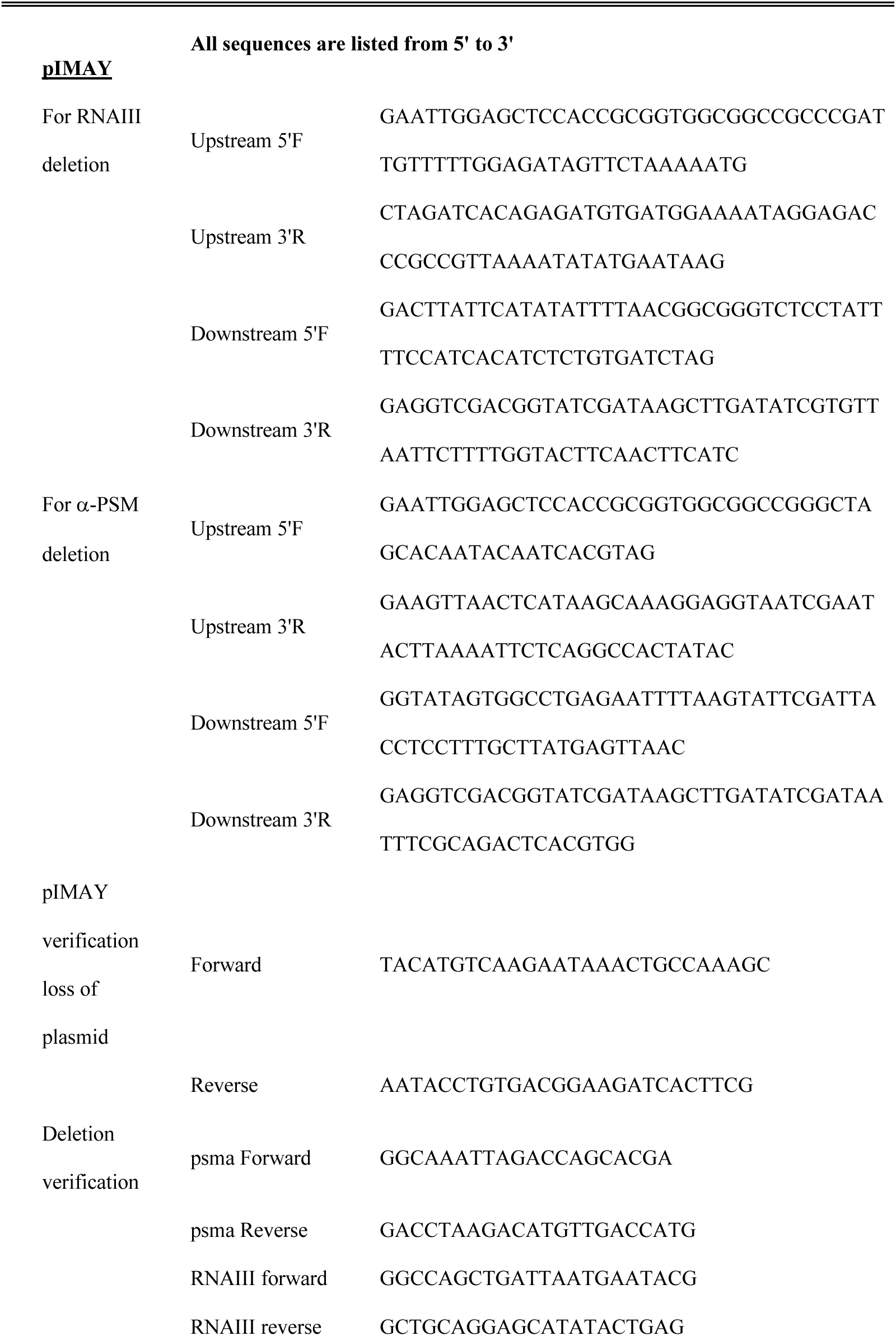
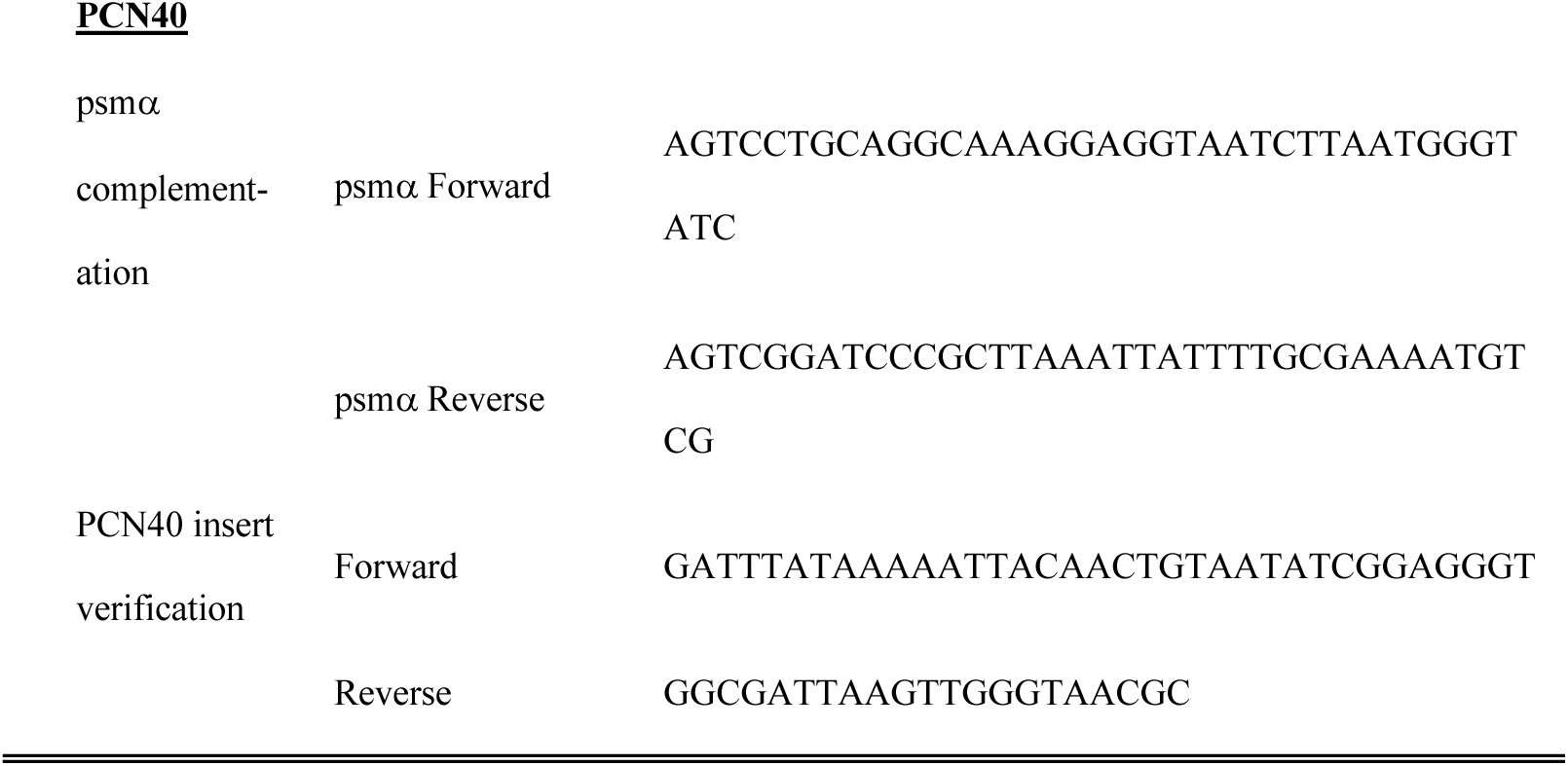
PCR primers used in this study.

### Gene ontology analysis and gene annotations

Gene Ontology (GO) analysis was performed using the ShinyGO version 0.85 database (Ge et al., 2020). KEGG and AureoWiki databases were used to as references for gene annotation (Fuchs et al., 2018; Kanehisa et al., 2021)

### Statistics

Statistical analyses were performed using GraphPad Prism version 8.0.2 for Windows, GraphPad Software, San Diego, California USA, www.graphpad.com

## Acknowledgements

The following reagents were provided by the Network on Antimicrobial Resistance in *Staphylococcus aureus* (NARSA) for distribution through BEI Resources, NIAID, NIH: Nebraska Transposon Mutant Library (NTML) Screening Array, NR-48501, *Escherichia coli* - *Staphylococcus aureus* Shuttle Vector pCN40, Recombinant in *Staphylococcus aureu*s, NR-46131, *Escherichia coli* - *Staphylococcus aureus* Shuttle Vector pCN48, Recombinant in *Escherichia coli*, NR-46143. *S. aureus* JE2 parent strain was a gift from Dr. Ken Bayles (Fey et al., 2013). pIMAY was a gift from Dr. Tim Foster (Addgene plasmid # 68939; http://n2t.net/addgene:68939; RRID:Addgene_68939) (Monk et al., 2012). *S. aureus* LAC strains and pTX vector were gifts from Dr. Michael Otto (Wang et al., 2007). We thank Dr. Ziqiang Guan for the lipidomic analysis of the JE2 *Δpsmα* and wild type strains. This work was supported by NSF Convergence RAISE (Research Advanced by Interdisciplinary Science and Engineering) award number 1931309, and Duke University Medical Center. Schematic illustrations were created with BioRender (https://BioRender.com).

## Supplemental Figures

**Supplemental Figure 1:**
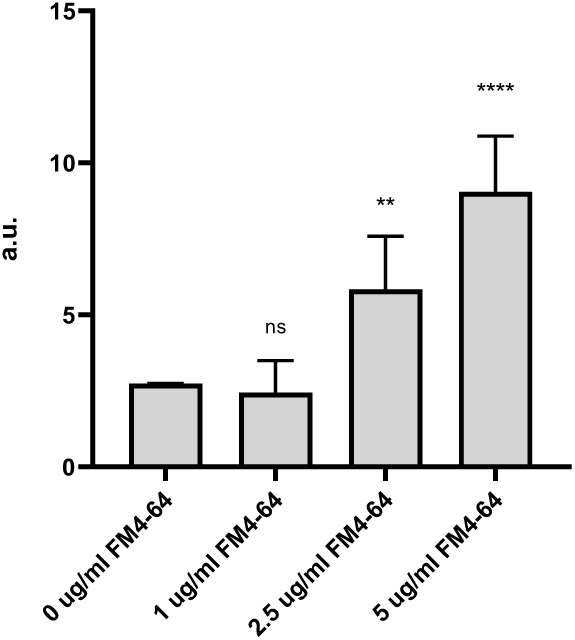
FM4-64 values of *S. aureus* JE2 cell free supernatants. Filtered cell free supernatants of wild type *S. aureus* JE2 grown in 96-well microtiter plates for 5 hours, incubated with 0, 1, 2.5, or 5 µg/ml of FM4-64. Values are presented as means from three independent experiments ± SD. Samples were compared using one-way ANOVA with Dunnett’s multiple comparison test *p< 0.05, **p< 0.01, ***p< 0.001, ****p< 0.0001

**Supplemental Figure 2:**
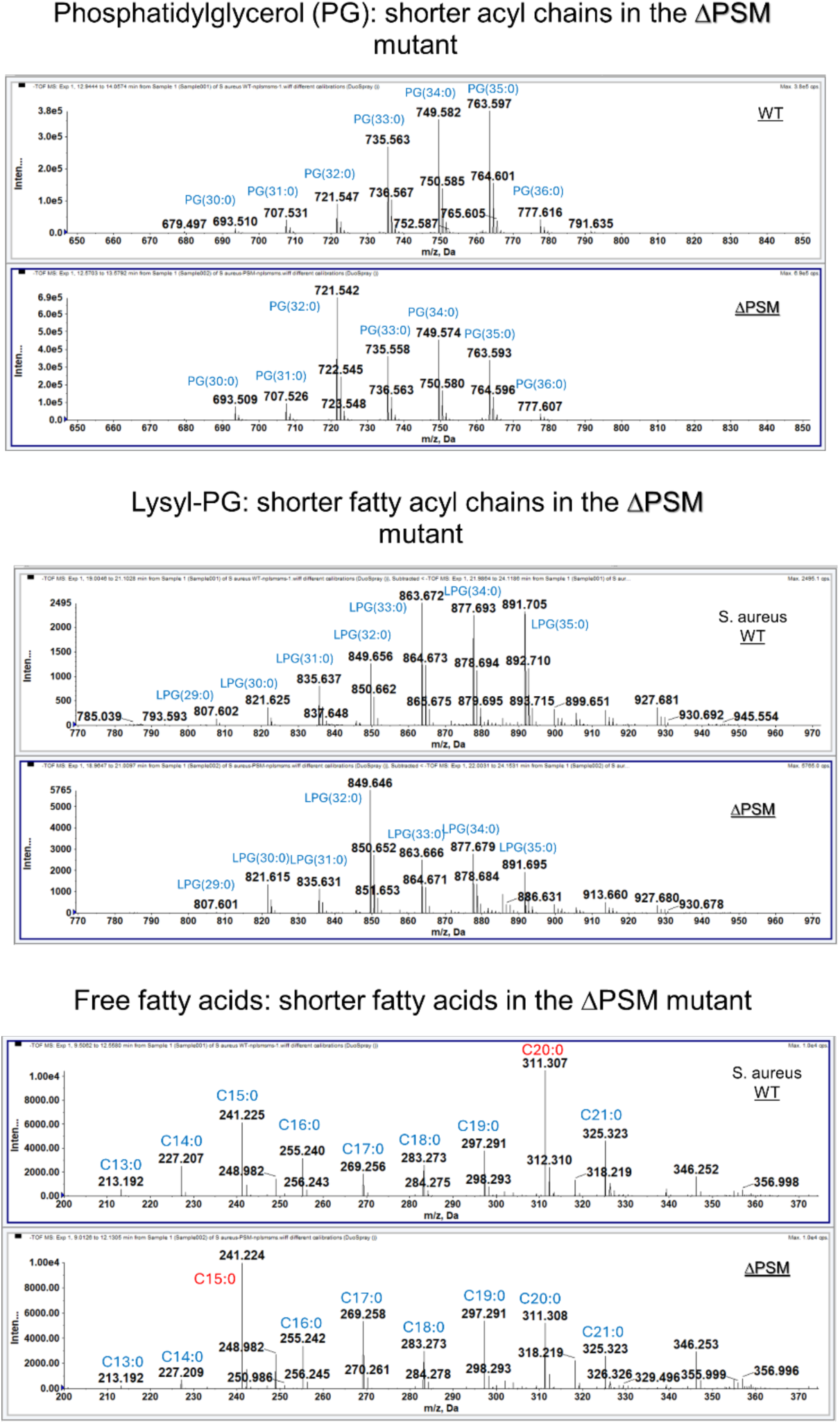
Lipidomic Analysis of JE2 wild type strain and JE2Δ*psmα.* Lipidomic analysis show that the JE2Δ*psmα* strain’s phospholipids contain shorter fatty acyl chains compared to the wild type JE2 strain. Analysis of free fatty acids show an increase in shorter fatty acids as well as branched chain fatty acids in the JE2Δ*psmα* strain

**Supplemental Figure 3:**
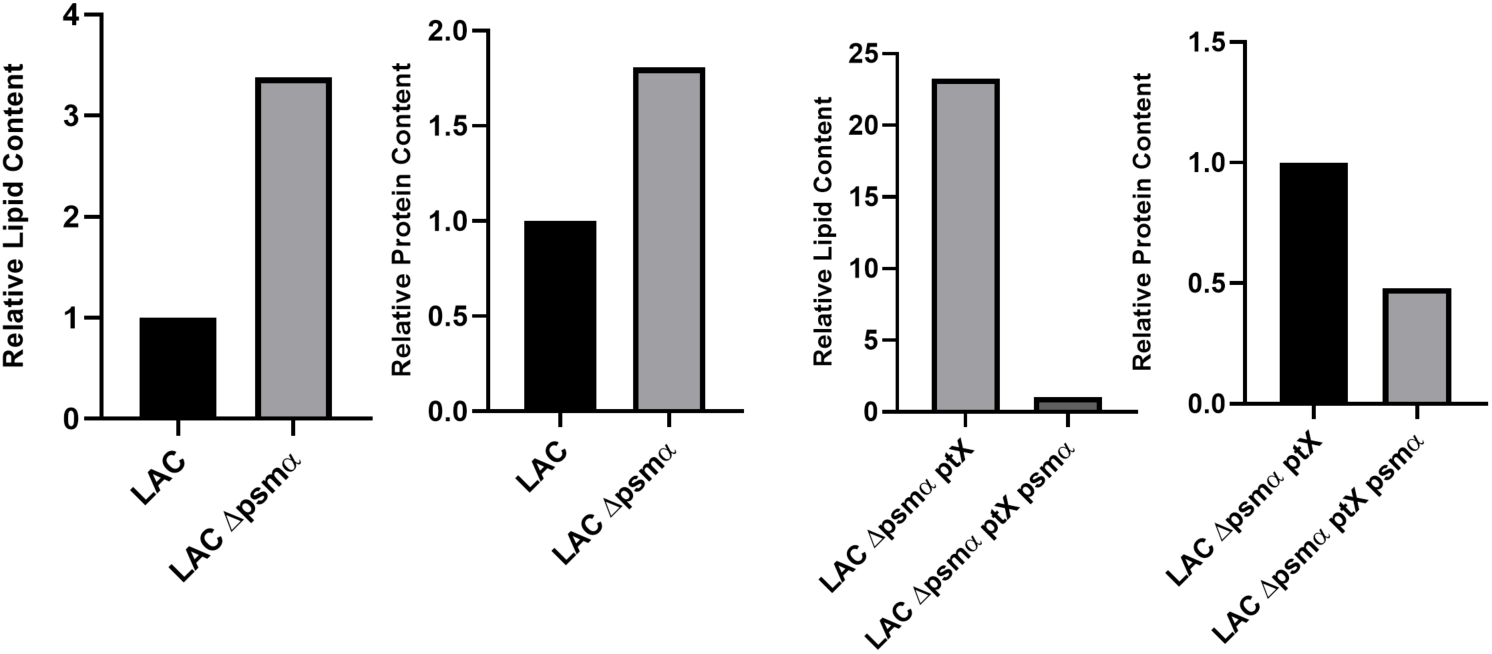
EVs from *S. aureus* LAC strains. The LAC Δ*psmα* (Wang et al., 2007) strain is hypervesiculating. Complementation with α-PSMs reduce vesiculation.

